# SIP-enabled multi-omics reveals soil microbiome responses to drought and rehydration

**DOI:** 10.64898/2026.03.30.715357

**Authors:** Tristan A. Caro, Juan Ibarra Arriaga, Eli Grossman, Aarushi Jhatro, Bryn Stewart, Alex L. Sessions, Smruthi Karthikeyan

## Abstract

The activity of the soil microbiome, and its balance of anabolic (organic C consuming) and catabolic (CO_2_-releasing) reactions, determines the magnitude and direction of soil carbon fluxes. Over half a century of research has revealed that soil water dynamics are key controllers of microbial activity. With increasing hydroclimate volatility expected across many regions of the Earth, there is a greater need to describe and quantify microbial responses to drought and rehydration cycles. In this study, we conducted rainfall exclusion experiments at two archetypical Mediterranean-type field sites. After rainfall exclusion and subsequent soil rehydration, we applied a SIP-enabled, multi-omics methodology to generate a multi-faceted case study of microbial growth, greenhouse gas fluxes, and the forms of carbon that drive both. Our results indicate that rehydration increases microbial anabolic processes by orders of magnitude, shifting cell generation times from years to days within just minutes. High-intensity drought increases the lag period before microbial growth resumes, but both stable-isotope probing and metagenomic inference agree that microbial communities exhibit greater capacity for rapid growth following drought stress. Furthermore, significant shifts in the soil metabolome are observed following drought and rehydration, implicating specific osmolytes as key to the microbial response and indicating metabolite diversity as a key modulator of microbiome functioning. Together, our results provide constraints on microbial activity rates in soil and mechanisms underpinning microbial responses to drought and rewetting. These findings motivate further research into microbial responses under increasingly volatile hydroclimate regimes and downstream contributions to the global carbon cycle.

**Significance Statement:** Soil is a major global store and source of carbon. The microbiome determine the fate of soil organic carbon, and the microbiome is ultimately controlled by soil water dynamics. Early, innovative experiments by H.F. Birch defined “The Birch Effect” – the observation that soils emit CO_2_ following drying and subsequent rehydration. However, it remains unclear when, and to what magnitude, soil microorganisms are actively growing following this rehydration, and what biological mechanisms explain the observed CO_2_ pulse. In this work, we apply an array of methodologies to address this question, describing rates of microbial growth during drought and rewetting. Our results provide crucial insights into how soil microbiomes will respond to increasing hydroclimate volatility across the globe.

## Introduction

Drying and rewetting is likely the most dramatic phenomenon regularly experienced by soil and its constituent microbiome. The addition of water rehydrates desiccated organisms, mobilizes organic and inorganic nutrients, and alters soil’s physical and hydrological characteristics (1). The response of soil microorganisms to drying and wetting has been a focus of soil microbiology for at least half a century, including the notable experiments of H.F. Birch who recorded patterns of CO_2_ flux following periods of drying/rewetting (2). Now termed “the Birch Effect,” the persistent observation of CO_2_ efflux from soils, with efflux rates modulated by soil drying history, carries with it critical implications for patterns of global soil carbon flow. In Mediterranean-type ecosystems, characterized by drought summers and winter rains, drying and rehydration is a critical modulator of the soil microbiome and C stocks (3). Increasing hydroclimate volatility, sometimes termed “hydroclimate whiplash,” is poised to increase the differential between dry and wet seasons in such ecosystems (4). California, for example, has experienced recent hydroclimate whiplash, and this trend is expected to increase over the next century due to anthropogenic climate change (5–7). The effects of these events on soil carbon and microbiome functioning remain difficult to anticipate, largely because our understanding of soil microbiome responses to drying/rehydration remains incomplete. Despite decades of study since Birch’s observations, major open questions persist regarding the timing, magnitude, and mechanisms underlying microbial growth during the drying/rewetting process, and how this parameter is affected by prolonged drought (1).

For microorganisms, growth depends on the extent to which a cell or community can assimilate and transform carbon, survive, acquire energy, and proliferate. Thus, growth is both an indicator of ecological and evolutionary phenomena (e.g., the extent to which an organism is adapted to its environment) as well as a reporter of geochemical transformation (e.g., carbon assimilation, respiration, etc.) (8). In the context of soil rehydration, the timing of microbial growth – defined here as anabolic processes that result in the production of biomass, remains unclear. Is growth tightly coupled to the flush of CO_2_ observed during the Birch Effect, or does it precede/follow it? What are growth rates during ambient dry conditions versus during rehydration events? Which forms of carbon fuel these anabolic processes? These questions are critically important to understand because the balance of microbial anabolic and catabolic processes dictates soil carbon dynamics. Recently, it has been demonstrated that microbial growth rate is the primary driver of soil organic carbon (9) and these processes are relevant at continental and global scales (9,10).

In this work, we sought to understand microbial growth responses to drought and rewetting by applying stable isotope probing (SIP) and multi-omics approaches. We conducted ^2^H-[vapor]lipid-SIP, ^18^O-DNA-SIP, gas flux measurements, metabolomics, and metagenomics at two case-study field sites during experimental rainfall reduction and rehydration treatments. Our quantitative, multiomics methodology reveals previously undescribed facets of soil microbiome function during periods of drought and rewetting. Specifically, we address the following open questions (i) what is the timing and magnitude of microbial growth following soil rehydration? (ii) how do these rates compare to ambient, dry-soil anabolic rates? (iii) how might different methods (lipid-SIP, DNA-SIP, genomic inferences) differ in their predictions of microbial activity? (iv) Which forms of carbon are most likely responsible for the microbial growth and flush of CO_2_ routinely observed following rewetting?

## Results and Discussion

### Experimental hydroclimate whiplash: two case-studies in California

We focused our study on two sites representing distinct California ecotypes of ecological and economic/agricultural significance in California: a grazed pasture and a coastal grassland (Fig. 1A). The pasture is located at Jalama Canyon Ranch, a farm near Lompoc, CA. This site hosts a sandy, hard-packed, clay-rich soil supporting primarily *Vulpia octoflora* (annual fescue), *Nassella pulchra* (purple needlegrass), *Medicago polymorpha* (Burr clover), and *Silybum marianum* (milk thistle). Goat and horse grazing occurs in the pasture throughout the year. The site typically experiences approximately 500 – 700 mm (∼20 – 27”) of annual precipitation. To induce a high-intensity drought treatment, a rainfall exclusion shelter (3 m x 3 m) of transparent polycarbonate sheeting was installed on Oct. 24, 2024. Vertical polypropylene sheeting was emplaced upslope of the shelter in order to reduce surface water flow along the hillslope into the shelter. This shelter successfully prevented precipitation infiltration, resulting in a volumetric water content (VWC) reduction in topsoil (0 – 10 cm) by the end of the experiment (**Fig. 2B**). This resulted in a significant reduction in plant cover (**Fig. 2A**). We refer to this pasture experiment as a “high intensity” drought condition throughout.

**Fig. 1.**
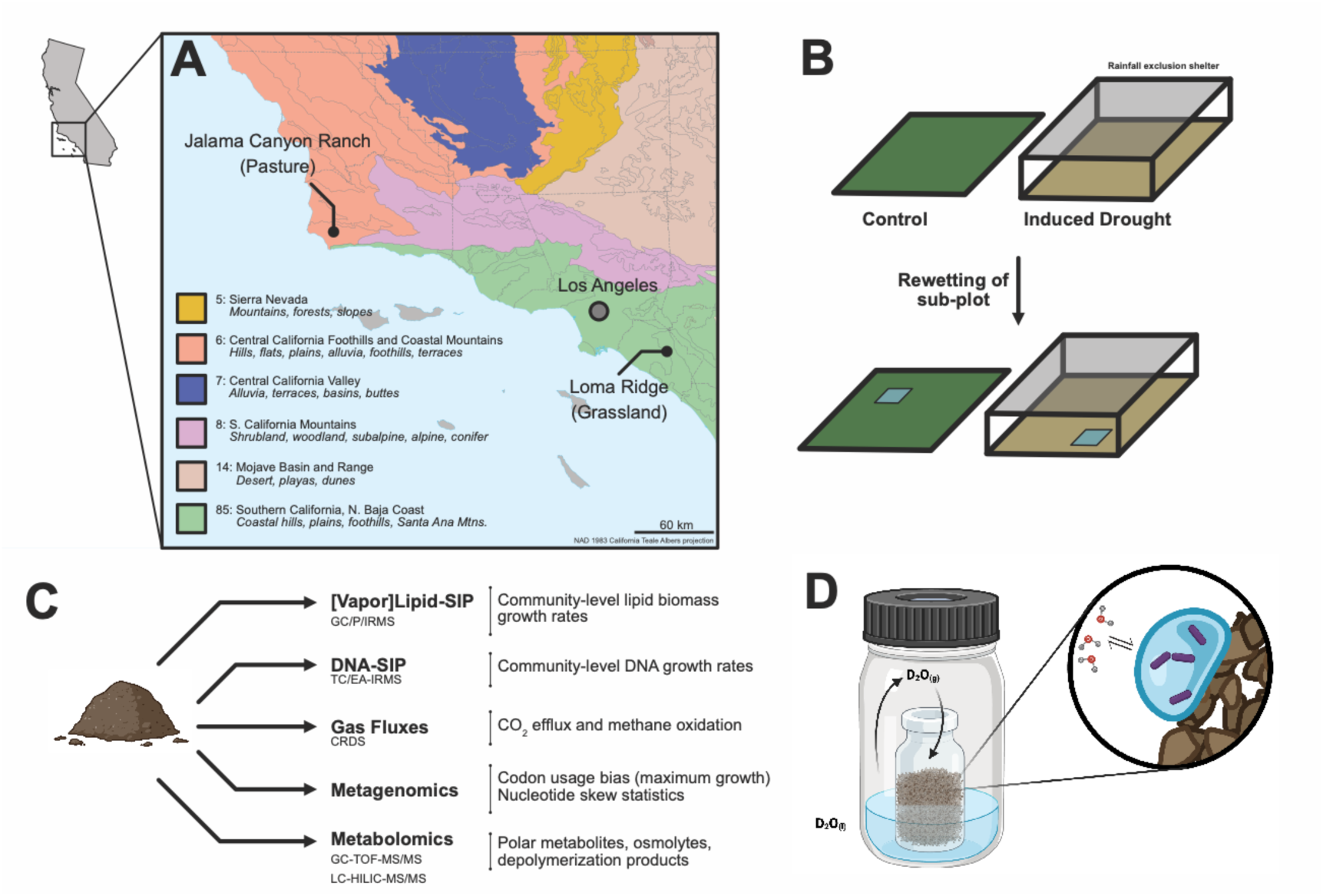
Experimental overview. **(A)** The locations of our two field sites, Jalama Canyon Ranch (pasture) and Loma Ridge (grassland) are plotted on a simplified ecotype map of California. Figure legend and color fill indicate ecoregion number and classification compiled as a multi-agency consensus, reported in Griffith et al (74). **(B)** Adjacent control and drought plots were sampled at the end of the rainfall exclusion experiment; rewetting of subplots was subsequently conducted. **(C)** Soils sampled from control and drought plots were analyzed by an array of techniques. Data outputs from each methodology are briefly described. **(D)** A schematic overview of the vapor/soil water equilibration method (25,29). Isotopically labeled water (D_2_O_(l)_) is passively equilibrated with water vapor in the incubation bottle headspace (^2^H_2_O_(g)_). This vapor equilibrates with soil pore water and provides an isotope probe to soil microorganisms contained within hydrated pores.

**Fig. 2.**
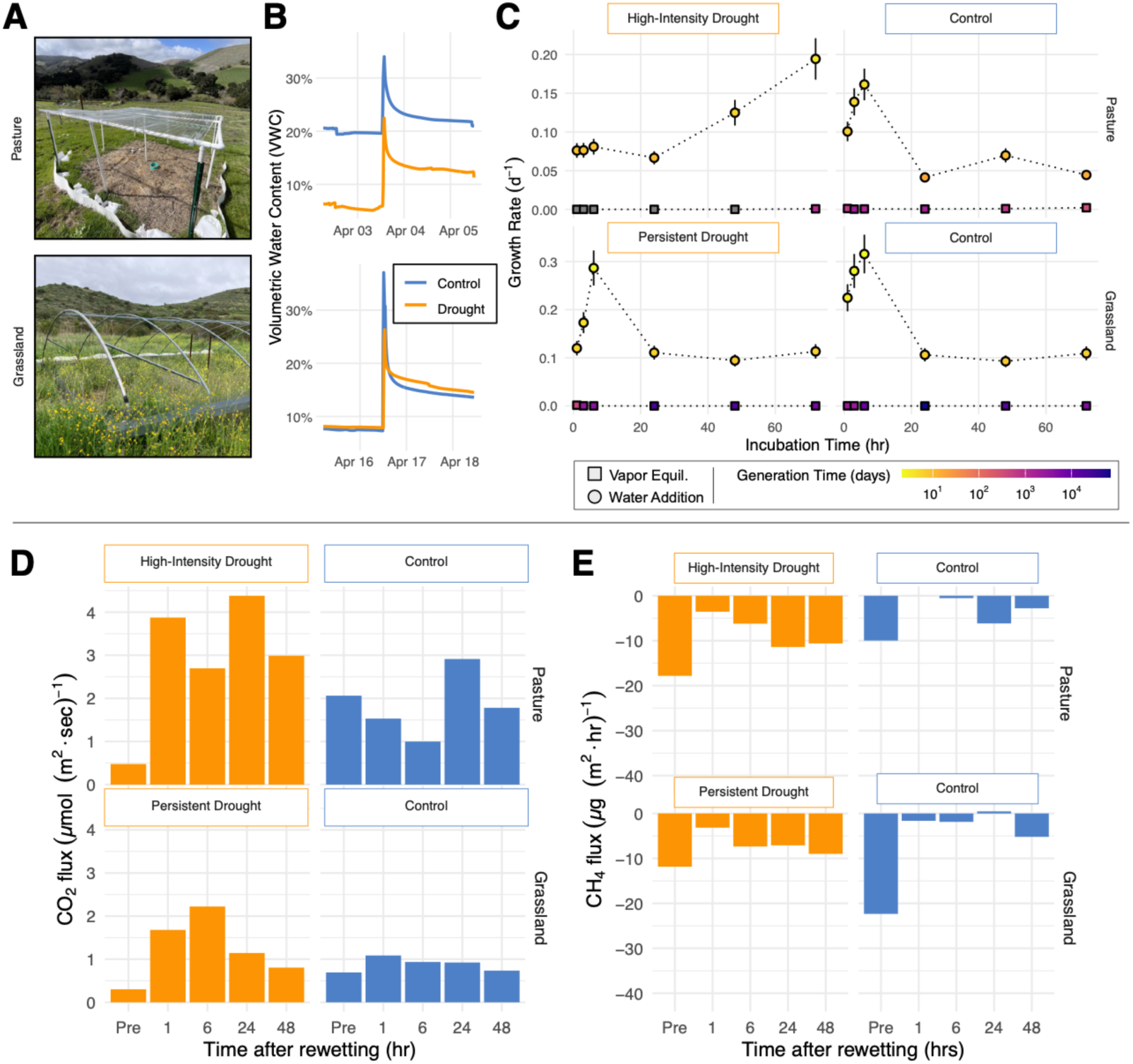
Growth and gas flux results. Note for all panels that the pasture (Jalama Canyon Ranch) and grassland (Loma Ridge) the drought experiments are referred to as “high intensity” and “persistent” drought, respectively. **(A)** Photographs from the two field sites: the pasture at Jalama Canyon Ranch (Lompoc, CA) and the grassland at Loma Ridge Global Change Experiment (Irvine, CA). **(B)** Volumetric water content (VWC) data immediately before, during, and after the rewetting experiment. Each plot is centered around the rewetting event, with an observed spike in VWC. Note that the 50% drought treatment in the Loma Ridge grassland site led to similar VWC prior to the rewetting experiment. **(C)** Lipid-SIP results are displayed; samples incubated using the vapor equilibration method are displayed as squares and those with traditional rewetting are displayed as circles. Error bars indicate the standard propagated error of measurements and growth rate calculations. Gray shapes indicate that growth was not detected. **(D)** CO_2_ flux measurements. **(E)** CH_4_ flux measurements. Negative values indicate loss (oxidation) of CH_4_. Note that gas flux measurements did not include a 72 hour timepoint due to field access constraints. For panels D and E, error bars are negligible and not shown.

The Loma Ridge Global Change Experiment, located in the Santa Ana mountains near Irvine, CA, hosts a persistent rainfall manipulation treatment in a hilled coastal grassland (**Fig. 1A**). The grassland consists primarily of *Bromus diandrus* (great brome grass)*, Avena fatua* (wild oat grass), *Dendra fasciculata* (clustered tarweed), and *Brassica nigra* (black mustard). The soil is a sandy loam and experiences approximately 325 mm (∼13”) of precipitation annually. Drought treatment has been underway at this site for 5 years (see *Materials and Methods*) and results in a 50% reduction of rainfall, rather than complete prohibition. Plant growth between the drought and control plots was visibly altered and reduced (11). Despite a clear reduction in rainfall intensity (water influx) long-term soil VWC measured at the site did not dramatically differ between the drought and control plots (**Fig. S1**). Greater water holding capacity by loam, and the partial rainfall reduction, likely explain the minimal effect on soil VWC. We refer to the Loma Ridge grassland experiment as a “persistent drought” treatment throughout, in contrast with the more extreme “high-intensity” drought experienced at the pasture site. We emphasize that these two field sites host significantly differing biomes, drought treatments, and land uses, and should be considered parallel experiments rather than contrasting treatments. Rainfall trends at each site for the 2024 – 2025 winter season are plotted in **Fig. S2.**

### Microbial growth responses to rewetting and decoupling from gas fluxes

In non-drought control plots at both sites, microbial growth rates reached their maximum rate 6 hours following rewetting (**Fig. 2A**). Intriguingly, the high intensity drought condition induced at the pasture site displays a substantial lag period before the maximum measured growth rate is reached at 72 hours following rewetting. In this case, microbial growth exhibits an extreme temporal decoupling from the onset of CO_2_ efflux (**Fig. 2D**), which begins within 1 hour following rewetting across all conditions. In contrast, the persistent drought condition, while still exhibiting a heightened CO_2_ efflux relative to the control, exhibits a tighter coupling of CO_2_ efflux and anabolic rates, which are both maximized at 6 hours post rewetting (**Fig. 2A, 2D**). Decoupling of growth and respiration in the high-intensity drought condition could be explained by a variety of factors. One hypothesis is that high-intensity drought drives surviving microorganisms to produce abundant osmolytes (compatible solutes) and that rapid respiration of these compounds and osmotic rebalancing is required before substantial anabolic activity can occur (12). It is also possible that this process co-occurs with the release of a large proportion of the microbial community from a dormant state, resulting in an increased lag period following the onset of more rapid growth. Indeed, our metagenomic dataset displays an increase of *Bacillus* in the extreme drought condition, indicating an enrichment in known spore-forming taxa (**Fig. S3)**. An alternate hypothesis explaining decoupling of CO_2_ efflux and growth is a contribution of relict CO_2_ to emissions. We define relict CO_2_ as CO_2(g)_ adsorbed to soil mineral/organic surfaces or otherwise trapped within soil pore matrices. Such gas is presumably of microbial origin but *not* produced via respiration concurrent with the measurement. Ostensibly, relict CO_2_ is released following rewetting due to alteration of soil structure and water-induced desorption (13,14). To assess this potential abiotic release mechanism, we conducted abiotic rewetting experiments of sterile, organic-clean silica sand (see *Materials and Methods*). These experiments indicate that abiotic CO_2_ release resulting from desorption and pore structure disruption occurs within approximately 9 minutes following rewetting (**Fig. S4**). Therefore, we contend that CO_2_ effluxes observed >1 hour following rewetting are unlikely to be the result of desorption of relict CO_2_.

Soil is a natural sink of atmospheric CH_4_ due to the activity of aerobic methanotrophs capable of oxidizing methane at atmospheric concentrations (15). Soil rehydration decreased methane oxidation rates across all conditions (**Fig. 2E**). Drought-stressed conditions recovered their CH_4_ oxidation capacity more rapidly. This result suggests that microbial CH_4_ oxidation is primarily limited by gas transport processes. In other words, in soils with greater volumetric water content (**Fig. 2B**), transport of sparingly-soluble gases such as CH_4_ may be limited by solubility and diffusion rates.

The ratio of soil CH_4_ influx to CO_2_ efflux can provide a rough estimate for the contribution of aerobic methanotrophy to soil CO_2_ release. This value ranges from 5.2 × 10^-8^ to 7.7 × 10^-4^ (mean = 1.1 × 10^-4^) across the rewetting event with the maximum observed under dry conditions (**Fig. 2**). Aerobic methanotrophy (CH_4_ + 2O_2_ → CO_2_ + 2H_2_O) in theory produces CO_2_ equimolar to consumed CH_4_, but a portion of this C is diverted to anabolic processes as formaldehyde into the RuMP cycle or formate into the serine cycle (16). Assuming a typical carbon conversion efficiency (CCE) of approximately 50% (16–19), we cautiously infer that approximately 0.006% of emitted CO_2_ from dry soils under ambient conditions, is the result of methanotrophy under ambient, aerated conditions. This fraction, again assuming a 50% CCE, would correspond to an approximate methanotrophic carbon assimilation rate of 167 pmol C (g · day)^-1^, a rate that roughly corresponds to the biomass of 1 to 5 methanotrophic cells’ worth of carbon fixed per g of soil on a daily basis, depending on cell-specific biomass (typically 30 – 120 pg C per cell) (20). Though the total population of methanotrophic organisms is not quantifiable, such an estimate suggests minimal contributions of CH_4_ to the SOC/biomass pool, despite soil oxidation still acting as a globally-relevant CH_4_ sink.

### Ambient rates of microbial growth in soil are extremely slow but rapidly up-shift

Microbial growth rate inference often relies upon the addition of an isotopically labeled compound (e.g., ^13^C-glucose, ^3^H-thymidine, etc.) in aqueous solution (21,22), or the application of an isotopically labeled-water (e.g., ^2^H_2_O, H_2_^18^O) (23,24). To assess the degree to which water addition stimulates microbial growth, we applied deuterated water in the gas phase (^2^H_2_O_(g)_), “vapor-lipid-SIP,” to estimate microbial anabolic rates under ambient, dry conditions. This method, first proposed by Canarini and colleagues (25) for tracing ^18^O-labeled water into DNA, passively equilibrates soil porewater with isotopically-labeled vapor in a hermetically-sealed vial. Isotopic composition of soil porewater is then estimated by tracking the isotopic distillation of the evaporating source water over time (**Fig. 1D, Fig. S5-S6**). Results from vapor-lipid-SIP indicate that soil microorganisms are growing at rates between 0.0001 – 0.0009 d^-1^ (mean = 0.0003 d^-1^) under ambient, dry conditions. These growth rate correspond to inferred doubling times on the order of years (**Fig. 2C**). Rehydration, as measured with traditional water-addition lipid-SIP, increases growth rates to approximately 0.1 d^-1^, an increase of 10^2^-to 10^3^-fold. Thus, ambient, dry rates of microbial growth are exceptionally slow, likely indicating the dominance of maintenance anabolism, complete stasis, and/or a lack of cell doubling. Interestingly, these ambient rates of growth are comparable to those previously measured by water-addition lipid-SIP in frozen subsurface permafrost environments (14). This observation suggests that extremely slow growth modes, traditionally ascribed to “extremophilic” taxa, are more cosmopolitan than previously thought (26,27).

Caro et al. had previously reported that DNA-and protein-amendment methods for measuring microbial anabolic rates in soil had likely overestimated *in situ* activity due to the introduction of C/N-rich compounds (e.g., glucose, thymidine, leucine, etc.), resulting in the “fertilization” of the soil microbiome (23). Our results reproduce this initial finding but further demonstrate that water addition itself, though nutritionally neutral, biases growth measurements of the soil microbiome, and that true *in situ* rates of growth are more reliably approximated by the vapor equilibration method. However, the vapor equilibration technique also includes caveats. Vapor equilibration necessarily increases the relative humidity (RH) of the air above the soil. Studies of soil microorganisms under desiccating conditions indicate that increasing RH itself, without direct application of water, can rescue cells from inactivity and dormancy (28). Thus, it is possible that humidity alone can revitalize components of the soil microbiome. Indeed, in our high-intensity drought experiment, we do not detect microbial growth via vapor-SIP until 72 hours following sealing of the vial, suggesting that growth is induced after 3 days following RH stimulation (**Fig. S7**). A potential mechanism explaining this result is the nucleation of isotopically-labeled vapor on microbial cell surfaces, thus promoting the expansion of new or pre-existing soil water films and pore waters. This hypothesis and the exact degree of isotopically labeled vapor infiltration into soil pore spaces, requires further experimentation to explore.

### DNA synthesis outpaces lipid synthesis

What can be learned from correlative measurement of DNA and lipid synthesis rates in soils? These rates inform our understanding of microbial physiology: biomass synthesis allocation, rates of replicative vs. non-replicative growth, maintenance/repair, etc. We sought to understand the relative rates of DNA and lipid synthesis in the soil microbiome. We initially considered three potential outcomes (**Fig. 3**). The first (i) is parity between lipid and DNA synthesis. In this case, DNA and lipid synthesis are tightly coupled following soil rewetting. Both lipid and DNA biomass synthesis are tied to growth and cell division, and imbalances are therefore minimized in order to optimize growth. (ii) Lipids are synthesized more rapidly than DNA. Non-replicative growth has been demonstrated in soils, especially in the context of storage compounds (29,30). Though we measure intact phospholipids, which are unlikely to be utilized in C storage compared to neutral lipids such as triglycerides, free fatty acids, etc., it is possible that lipid biomass will be turned over more rapidly due to membrane remodeling, repair, or growth in the absence of DNA replication. Indeed, if microbial growth is slow, it is possible that lipid synthesis may occur prior to the onset of the C-period and thus, DNA synthesis may not be captured on short timescales. However, our results align squarely with the third possibility (iii), wherein DNA synthesis routinely outpaces lipid synthesis. In our samples, this excess activity is by a factor of 2-10X (mean: 3.41X), depending on site and condition (**Fig. 3B**), and this disparity increases with time after rewetting. This result may be explained by a variety of DNA synthesis processes in excess of genome replication e.g., DNA repair, DNA phage replication, MGE synthesis, etc. However, we consider one possibility to be particularly intriguing: ploidy changes resulting from observed growth shifts. It is known that bacteria in culture may change their ploidy levels (replicate genome copies) to accommodate more rapid growth (31) and polyploidy may result from cell-cycle dysregulation under nutrient-replete conditions (32). However, the ploidy levels of bacteria in environmental samples are almost entirely unexplored (33). Whether bacteria in natural systems, experiencing growth rates far slower than cultured environments, are routinely experiencing polyploidy, remains an open question. Further experimentation is required to adequately substantiate the hypothesis that environmental microbiomes experience ploidy shifts with increased growth rate.

**Fig. 3.**
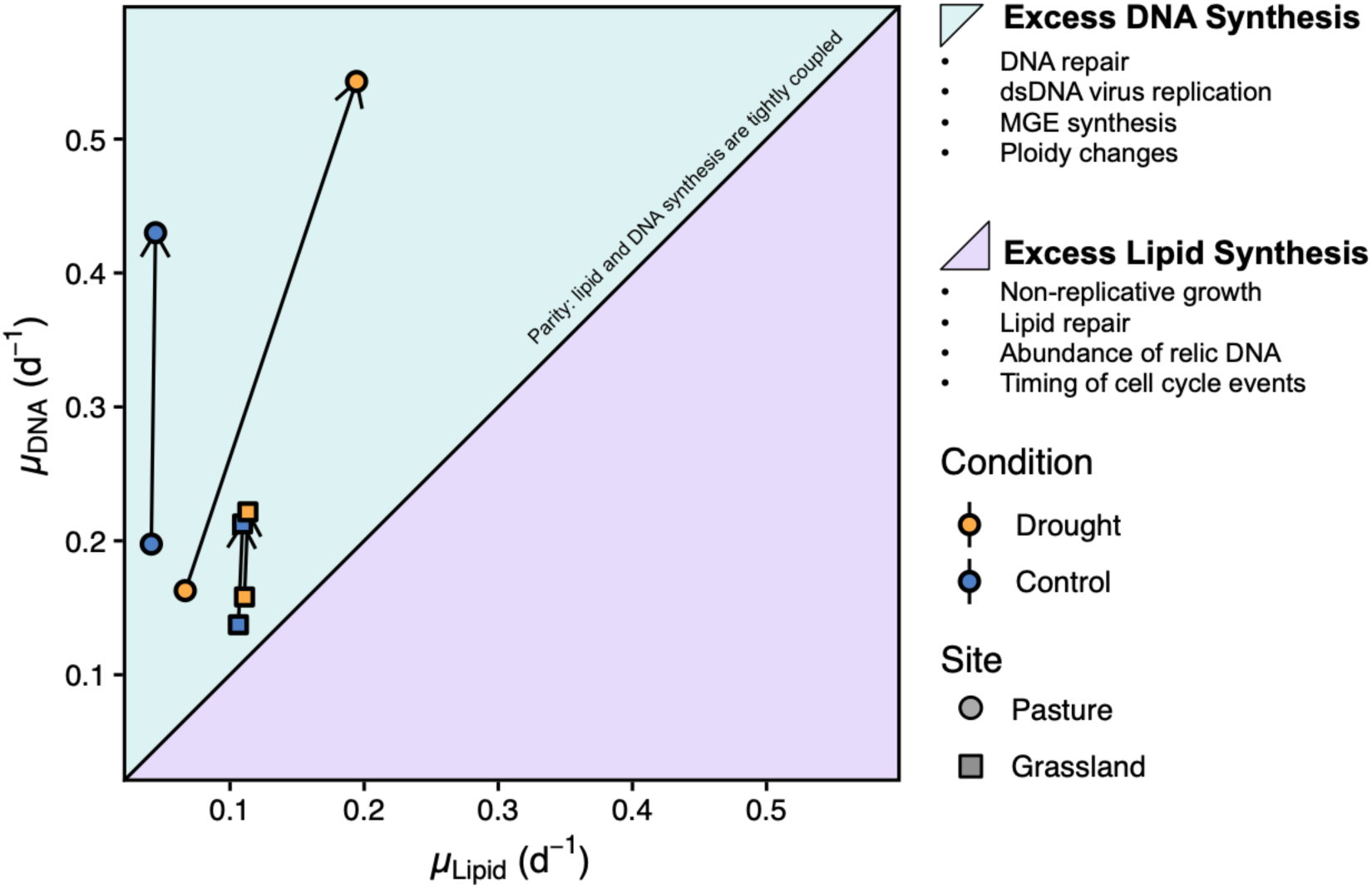
Comparison of DNA and lipid synthesis rates derived from comparable SIP datasets. Differences in lipid-and DNA-synthesis rates, calculated as relative growth rate derived from DNA-SIP (µ_DNA_) and lipid-SIP (µ_lipid_). The purple area indicates lipid synthesis in excess of DNA synthesis; the teal area indicates DNA synthesis in excess of lipid synthesis. Arrows mark the progression of time, with the arrow tail denoting the 24 hour SIP incubations, and the arrowhead noting 72 hours following rewetting.

An alternate possibility explaining this result is purely computational and concerns the assumed “water assimilation efficiency” (a_w_) parameters of each method. For both lipid-SIP and DNA-SIP, one must estimate the fraction of lipid hydrogen and DNA oxygen, respectively, that are sourced from H_2_O. This a_w_ term introduces great uncertainty into calculations because it is a direct scaling factor on estimate growth rates (34,35). For lipid-SIP in soil, lipid H a_w_ has been assumed to be ∼0.7. For DNA-SIP, a_w_ of DNA O is thought to be much lower, ∼0.3 (23,24). In order to achieve parity between the two methods DNA O a_w_ would have to be increased substantially **(Fig. S8)**. Further research estimating potential ranges of DNA O a_w_ in soils is required to further constrain this term and refine growth rate metrics.

Whether driven by ploidy shifts, MGEs, viral blooms, methodological biases, or other unknown mechanisms, a critical implication of differential DNA and lipid synthesis rates concerns carbon biogeochemistry. Global-scale estimates of carbon-use efficiency, microbial growth, SOC stores, and SOC dynamics, are modeled using ^18^O-DNA-SIP-derived to calculate a relative rate of microbial growth (d^-1^) (9,36–38). Relative growth rates derived from DNA replication are multiplied by estimates of microbial biomass C to estimate absolute growth rates of microbial biomass C (e.g., µg C d^-1^) (8). Previous work has indicated that C/N labeling approaches over-estimate microbial growth rates in soil (23,37,38); our work suggests that ^18^O-DNA-SIP does as well. Though counter-intuitive, the synthesis of DNA within a microbial cell may be decoupled from its rate of biomass production. Though [vapor]-lipid-SIP may be instrument intensive, measuring the production of new lipid biomass may provide a more appropriate proxy for microbial biomass C production because membrane growth is (i) sensitively measured, even in the absence of cell fission and (ii) is more directly related to microbial biomass carbon. Lipid-SIP provides extremely limited taxonomic information (23,29), and so the ideal approach is multi-modal, with lipid-SIP providing insight into anabolic rate, and DNA SIP identifying relevant taxa/genes implicated in microbiome function (8).

### Genomic indicators of microbial growth potential shift under drought conditions

Metagenomes may hold clues regarding microorganisms’ growth capacity. Vieira-Silva & Rocha (39) demonstrated that, across an array of genomic metrics (e.g., rRNA copy number, tRNA copy number, etc.), high codon usage bias (CUB), the tendency of organisms to bias the use of specific codons, is strongly associated with an organism’s maximum growth rate. Later work by Weissman et al. (40,41) improved the predictive capacity of CUB statistics and expanded their applicability to metagenomic datasets. We sought to build upon this prior work and evaluate the degree of correspondence between maximum predicted growth rate (inferred from CUB statistics) and empirically-measured growth rate (measured via lipid-SIP). In other words, does *predicted* growth potential from genomes, at the community level, correspond with *actualized* growth rates? To answer this question, we conducted standard metagenomic assembly and binning (see *Materials and Methods*) on samples collected in parallel to our water-addition lipid-SIP dataset, resulting in the assembly of 73 metagenome-assembled genomes (MAGs). We estimated codon-usage bias (CUB) patterns of resulting MAGs. CUB-derived estimates of maximum growth rates indicate that drought-stressed microbiomes at our sites exhibit significantly greater capacity for fast growth than those in corresponding control plots (*P = 0.013, Tukey’s*) (**Fig. 4A**). This trend holds at both sites when examined individually (**Fig. S9**). This is intriguing in light of our lipid-SIP dataset, which indicates a longer lag period for microorganisms under high intensity drought stress. However, the maximum growth rates observed via lipid-SIP are in the high-intensity drought condition, with mean microbial generation times on the order of approximately 4 days (**Fig. 2A**). Under high-intensity drought conditions, this capacity for rapid growth is actualized following a relatively sluggish lag period. Thus, to a first order, growth *capacity* does not directly translate to growth *actualization* in the short term but may do so given time or more favorable conditions following perturbation. This result supports prior observations that greater codon usage bias is associated with taxa that more rapidly synthesize DNA following soil rewetting (42). Also in concordance with previous work, we note an apparent taxonomic signal of CUB-predicted growth potential: MAGs belonging to *Bacillota* exhibit the shortest minimum doubling times; *Verrucomibrobiota* exhibit the longest, though within-phylum variation can be substantial (**Fig. 4B**) (8,39–41).

**Fig. 4.**
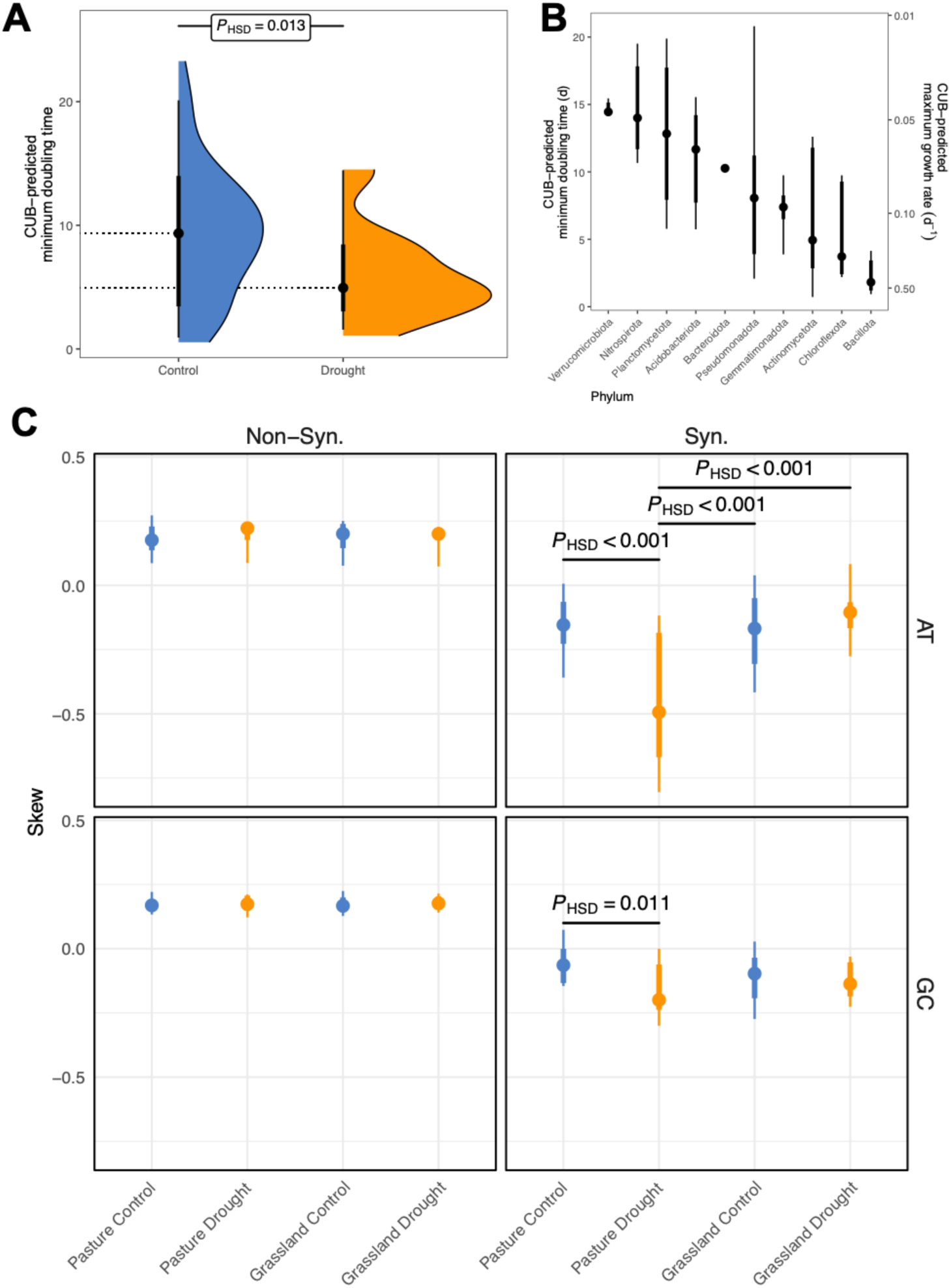
Genomic parameters influencing growth responses. Distributions of parameters calculated from MAGs. Points indicate median, thick and thin bars indicate 66% and 95% CI, respectively. Significance indicated is Tukey’s honestly significant difference (P_HSD_) test. **(A)** Comparison of CUB-predicted minimum doubling times between drought and control plots. Dotted lines emphasize median value. Slabs denote kernel density. MAGs from both study sites are aggregated. **(B)** Comparison of CUB-predicted minimum doubling times of all MAGs assembled in this study, annotated at the phylum level. Right y-axis indicates the transformation of minimum doubling time to maximum growth rate. **(C)** Nucleotide skew statistics, either A→ T or G→C at non-synonymous (non-syn), synonymous (syn) coding sites.

We also assessed nucleotide skew in our metagenomic dataset. Nucleotide skew is defined as the tendency of microorganisms to preferentially utilize A over T and G over C in their genome. This parameter has been associated with more rapid transcription, as certain nucleotides are more energetically costly to produce: A > T, G > C. (43). Transcription-related selection is thus thought to favor the energetically “cheaper” nucleotides U and C. Nucleotide skew at synonymous coding sites of ribosomal proteins has been associated with increased transcription efficiency in soil (42). In our dataset, high-intensity drought-associated MAGs exhibit significant synonymous AT and GC nucleotide skew (p<0.001, p=0.011, respectively, Tukey’s), suggesting that organisms under high-intensity drought, in addition to exhibiting more rapid growth (**Fig. 4A**), also streamline their ribosomal protein genes to favor inexpensive nucleotides (**Fig. 4C**). Nucleotide skew is not exhibited at non-synonymous sites of ribosomal protein sequences (**Fig. 4C**), suggesting that this is a translation-level selection process, rather than a force acting on genome encoding itself. In addition to ribosomal protein genes, explored by previous studies (42,43), we also examined nucleotide skew across the full MAGs, finding significant nucleotide skew in a variety of cellular functions. Binning genes across COG categories, significant GC skew was observed across all categories (p<0.001); significant AT skew observed across all categories (p<0.01) except for one (**Fig. S10-S11)**. This finding suggests that nucleotide skew is exhibited across the entire genome in organisms selected for by high-intensity drought. These organisms appear to be survivalist/opportunists: organisms that can survive desiccating conditions but also exhibit the capacity and adaptations for rapid proliferation upon encountering more favorable conditions. Interestingly, within the YAS (growth vs. assimilation vs. survival) ecological tradeoff framework, such organisms appear to strongly associate with both survival and growth axes (Y-S) (44,45). One might hypothesize that such taxa may be deficient in their degradative (“A”) capacity – that is, limited in the range of C that they can transform and assimilate. This phenomenon is suggested by our metabolomic data, discussed below.

### Drought and rehydration modulate the metabolome

We observe metabolomic shifts of soil both accumulated over the months-long drought period as well as during the short-term response to rewetting. Drought conditions (both high-intensity and persistent) significantly shift the bulk metabolome of soil (**Fig. 5A**). Interestingly, drought results in greater compound accumulation than loss. This likely indicates multiple co-occurring processes including (i) accumulation of primary metabolites resulting from a reduced microbial capacity for OM degradation; (ii) accumulation of polar metabolites that cannot be transported under low water conditions (iii) microbial production of osmolytes during drought stress. Of the many polar metabolites that may have osmotic protection roles, we conservatively identify only a few that have well-characterized physiological roles in osmotic shock protection. These known osmolytes include: ectoine, 5-hydroxyectoine, trimethylamine N-oxide (TMAO), betaine, proline-betaines, trehalose, and mannitol (46–51). All of these osmolytes increase in concentration under drought conditions, with the majority being significantly accumulated (p < 0.05) (**Fig. 5B**). Indeed, the metabolome of both sites is distinct, both with and without drought considered (**Fig. 5C**). Principal component (PC) analysis of the metabolome provides clues (**Fig. S12**). 6 of the top 10 loadings of PC1 are compounds homostachydrine, N-methylproline, 5-aminovaleric acid, sulfoacetic acid, indole-6-carboxaldehyde, and beta-sitosteryl sulfate. 5-aminovaleric acid (5-AVA) Homostachydrine and N-methyl proline are both quaternary nitrogen compatible solutes, sharing their functional class with betaine and protein betaine. 5-aminovaleric acid is an end-product of proline catabolism, indicating an active proline cycling response under stress. Thus, PC1 is primarily an N-methyl osmolyte and metabolism axis and drought-induced nitrogenous osmolyte accumulation is the dominant source of variance in the dataset. PC2 is less clear-cut. We tentatively suggest that PC2 represents a broad site effect with separation originating from plant inputs differentiating a loamy grassland from a clay-rich pasture (52,53).

**Fig. 5.**
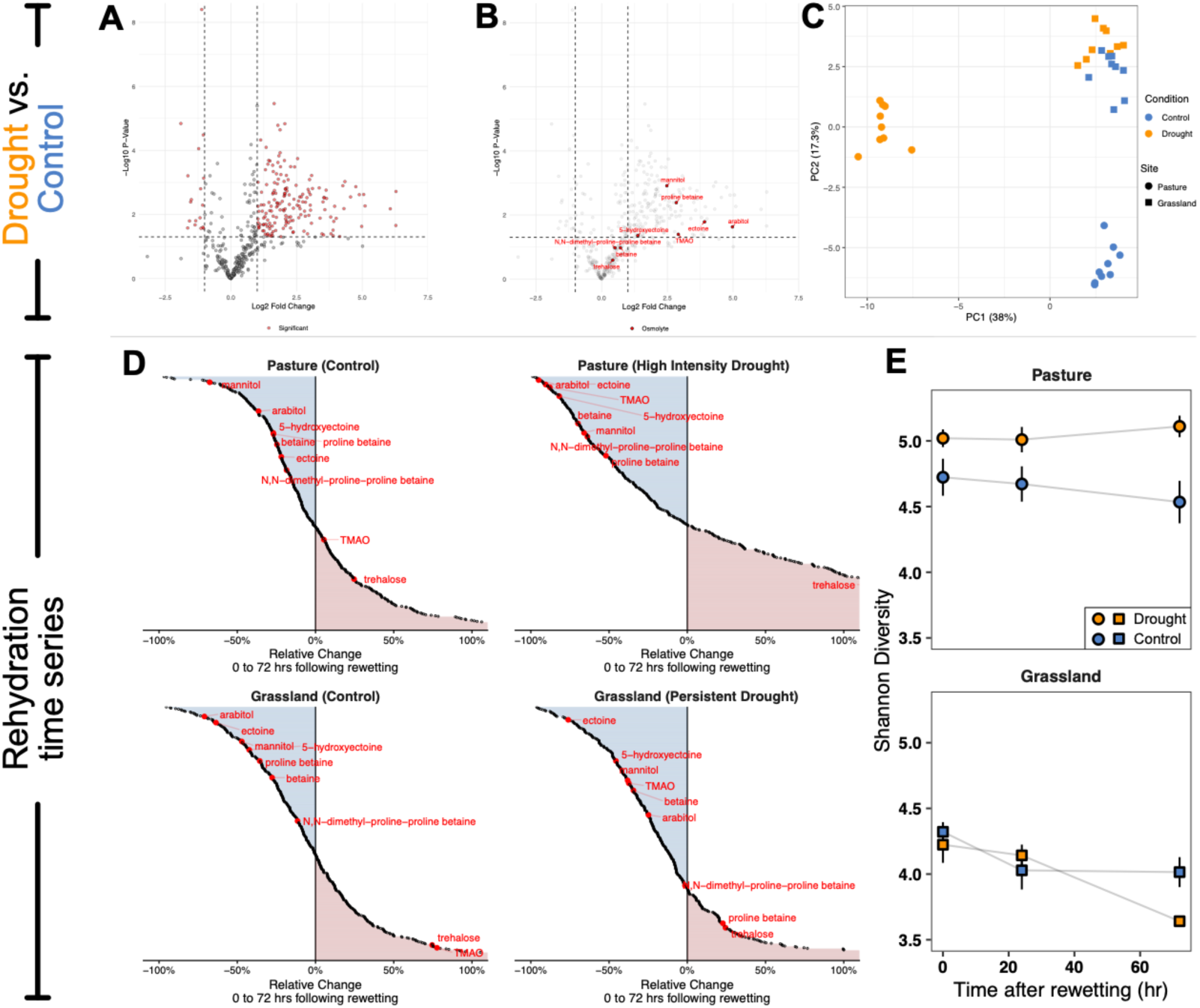
Metabolomics results. **(A)** Volcano plot of combined GC and LC tandem mass spectrometry datasets. Red points indicate significance, defined as > 1 Log2 fold change and p < 0.05). Fold change denotes increase or decrease in the drought conditions compared to control. **(B)** Subset of data in Panel A with known osmolytes indicated. **(C)** Prinicpal component plot of metabolomics dataset. Each point represents individual soil replacte samples from each experimental treatment. Treatments are distinguished by color and shape. **(D)** Plots display fractional relative change in metabolite abundance across the rehydration experiment. Known osmolytes are indicated in red. (**E**) Plots display metabolite alpha diversity (Shannon’s) within each field site. Error bars indicate standard error of experimental triplicates. Note that pasture and grassland drought conditions are “high-intensity” and “persistent” drought, respectively (Fig. 1C).

A persistent mystery in soil microbiology is the exact chemical origin of the flush of CO_2_ observed following soil rewetting (1,46). Does this CO_2_ come from the depolymerization of newly mobilized soil organic carbon, from the rapid respiration of intracellular compatible solutes (osmolytes), or both? In previous work on experimentally-dried soils, Warren (46) suggested that both osmolyte compounds and degradation products of OM accumulate and may contribute to the flush of observed CO_2_. Our results support this. Our metabolomic dataset indicates a greater concentration of metabolites (rather than loss) under dry conditions (**Fig. 5A**). Expanding on earlier experiments by Warren and others, we find that rehydration of soil leads to a rapid loss of many polar metabolites, with osmolytes being among those most rapidly lost (1,12,46,50,54). We interpret this widespread loss of metabolite abundance as a broad stimulation of carbon degradation, cycling, and respiration processes during soil rehydration. We were surprised to observe that trehalose consistently *accumulates* in soils following rewetting. This appears to be in conflict with prior studies that implicate trehalose specifically as a primary respiratory substrate for microorganisms responding to rewetting events (54). Though our polar metabolite extraction is intended to lyse cells (thus our analysis is intended to include both intracellular and extracellular osmolytes), we suspect that this observation may be the result of trehalose being ejected out of cells into extracellular space, rather than directly respired.

Metabolome alpha diversity (Shannon’s) was elevated under high-intensity drought conditions (pasture) **(Fig. 5D)**. No significant difference was observed under persistent drought conditions (grassland) between drought and control conditions (**Fig. 5D**). Following rewetting, decreases in metabolome alpha diversity were observed in persistent drought conditions (grassland) and less pronounced, but still significant decreases were observed in the pasture control site (**Fig. 5D**). No clear loss of metabolite alpha diversity is observed following rewetting under high-intensity drought conditions. Previous model-based analyses by Weverka et al. predicted an inverse relationship between metabolite alpha diversity (“chemodiversity”) and microbial carbon assimilation processes (55): our study provides empirical evidence for this relationship. Increased chemodiversity is thought to suppress assimilatory processes because of (i) increasing the pool of inaccessible or poorly-accessible substrates and (ii) increasing the enzyme synthesis costs associated with accessing these pools. Thus, increased chemodiversity under drought conditions likely reflects a decreased capacity of the soil microbiome to degrade soluble organic matter over time, likely leading to the accumulation of diverse organic substrates (56–58). Our metabolomic dataset is therefore consistent with the YAS framework, wherein organisms in high-intensity drought conditions, which exhibit substantial growth and survival capacities (as defined by their high codon usage biases and genome-wide nucleotide skew) experience “trade-offs” and sacrifice resource acquisition traits (44,45).

### Conclusions and Implications

This study presents a multi-modal, case study analysis of microbial growth responses to drought and rewetting at two archetypical Mediterranean-type sites in California. Increasing hydroclimate volatility is expected in Mediterranean-type ecosystems like California but changes to soil microbiomes and subsequent soil carbon cycling dynamics are not accounted for in these forecasts (4,5,59). In this work, we illustrate several key characteristics that may be expected when drought-stressed soils experience precipitation (summarized in **Fig. 6**). First, rapid and sustained fluxes of CO_2_ are expected as accumulated SOM and osmolytes are rapidly respired. As initially observed by Birch, these fluxes are greater following more extreme drying events (2,3). We propose that this is the result of a stressed, dysbiotic soil microbiome exhibiting a reduced anabolic efficiency (manifested as a lowered soil carbon-use efficiency) associated with traits related to survival and growth (as defined by the YAS framework) (44,45). Simultaneously, microbiomes in saturated soils rapidly lose their capacity for aerobic methanotrophy, as gas diffusion, especially of relatively insoluble CH_4_, is substantially hindered. Second, microorganisms that were largely dormant or operating under maintenance regimes revitalize and increase their growth rates by 100 - 1000s fold. This growth up-shift is largely driven by organisms adapted for rapid growth (as inferred from CUB patterns) and possibly induces ploidy shifts to accommodate the need for rapid DNA synthesis. Finally, growth is supported both by osmolyte respiration and the degradation of newly-mobilized SOM. Drought-stressed soils exhibit an increase in chemodiversity, likely driven by decreased degradation capacity of the attendant soil microbiome. This elevated chemodiversity may disrupt normal degradative processes. Taken together, our results suggest that an increasingly volatile hydroclimate in Mediterranean-type ecosystems may induce severe soil carbon cycling dysregulation that directly stems from altered soil microbial growth traits and rates. Increasingly dysbiotic soil microbiomes, characterized by stress, growth inefficiency, and traits favoring survival and weed-like growth over persistent carbon degradation could potentially lead to net C loss from soils over seasonal and annual timescales. Such losses of C from dry ecosystems, driven by precipitation, have already been observed by remote sensing on the continental scales (10). Our case studies motivate the need for urgent follow-up, where sensitive multi-omic SIP techniques are deployed in large field-based trials across a greater diversity of soil environments and future hydroclimate conditions. Such studies are required to clarify and quantify the microbially-mediated mechanisms underpinning potential soil-climate feedbacks at regional and global scales.

**Fig. 6.**
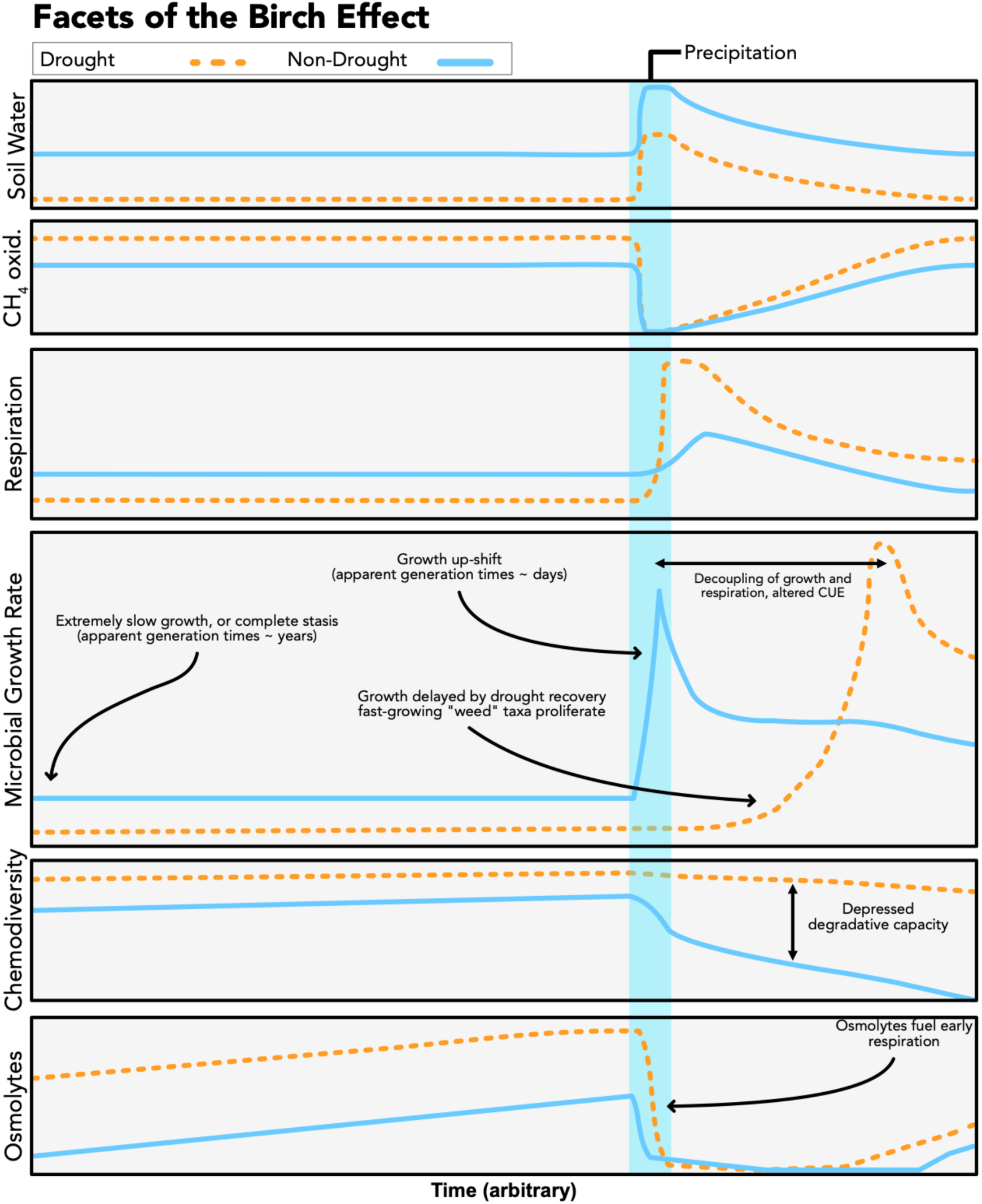
Facets of the Birch Effect. In this figure, we synthesize suspected facets of the Birch effect, both those previously characterized and indicated by our datasets. The vertical axes are relative and arbitrary, intended to serve as a visual aid. The x-axis is generalized, meant to represent broadly periods prior to, during, and following soil rehydration.

## Materials and Methods

### Field treatments and soil analysis

Rainfall manipulation experiments consisted of a “conditioning period” during which soils were subject to drought or control treatments. At Jalama Canyon Ranch (pasture) this period comprises Oct. 24, 2024, to April 5, 2025. Drought treatments at Loma Ridge began in February 2007. Each drought plot is equipped with a retractable canopy that is closed for selected rainstorms with the intent to reduce precipitation by approximately 50% compared to control plots. In October 2020, the Silverado Wildfire burned through Loma Ridge, destroying drought treatment structures. The structures were replaced in January 2021, at which point the experiment resumed (60). Soil volumetric water content and temperature were continuously monitored at both sites using TOMST TMS sensors emplaced at 10, 20, and 30 cm depth.

After each conditioning period, precipitation was simulated via the addition of 20 L of water over a 1m^2^ subset of the plot area at a rate of 0.33 L/min, resulting in hydration in-excess of field capacity. Soil CO_2_ and CH_4_ fluxes were measured using a LI-7810 CH_4_/CO_2_/H_2_O trace gas analyzer (LI-COR) using a temperature-monitored static chamber. Gas fluxes were determined by the increase or decrease of CO_2_/CH_4_ within a static chamber over a 15 minute monitoring period.

Precipitation data at JCR has been collected since October 2022 via a Davis GrowWeather (Vantage Pro2) weather station. Precipitation data at Loma Ridge (Hicks Canyon) has been collected by the Orange County Public Works department since 2007 and is publicly accessible at http://hydstra.ocpublicworks.com/web.htm. Precipitation data for both sites is displayed in **Fig. S1**.

Accurate estimates of soil grain size distributions are required for VWC data calibration. For soil grain size determination, 100 mg of each soil was decarbonated with 10 mL of 1M HCl, centrifuged at 3000G for 15 minutes, decanted, and rinsed 3X with 30 mL deionized H_2_O. Organics were removed by incubation with 5 mL of 30% H_2_O_2_ at 60°C for 30 minutes. Samples were centrifuged, H_2_O_2_ decanted, then 10 mL sodium metaphosphate solution (2.5 g/L) was added to each sample and sonicated for 10 minutes. Samples were then introduced to a Mastersizer 3000+ Pro for grain size determination.

### Lipid Stable Isotope Probing

Lipid-SIP was carried out using deuterated water in either a liquid or vapor form. Water-addition lipid-SIP was conducted following the procedure described in Caro et al. (23): 6 g of sieved soil was weighed into a plastic 50 mL centrifuge tube. 6 mL of 0.5 at. % D_2_O was added dropwise to the soil and allowed to incubate for 0, 1, 3, 6, 24, 48, or 72 hours. At the end of the incubation, the soil was centrifuged and water removed, an aliquot taken for water isotope analysis to account for dilution by soil water. Soil was flash-frozen by plunging in a dry-ice ethanol bath to halt microbial activity. Soil was lyophilized for 48 hr prior to extraction. Vapor-phase lipid-SIP was conducted similarly to the procedure described by Canarini et al. (29). In brief, 6 g of sieved soil was placed in a glass 20 mL vial and lowered into a 250 mL media bottle containing 6 mL 20 at. % D_2_O. The media bottle was sealed with a butyl stopper. Incubations were terminated after 1, 3, 6, 24, 48, and 72 hours, and D_2_O at the bottom of the vial sampled for water isotope analyses.

Intact polar lipids were extracted following the method defined by Buyer & Sasser (61). In brief, 5 g of sieved, lyophilized soil was placed in a glass centrifuge tube with PTFE-lined caps. 50 µg of 23:0 phosphatidylcholine was added as an internal standard. The following extraction mixes were applied in sequence: (A) 4 mL of 50 mM K_2_HPO_4_ buffer, (B) 10 mL methanol (MeOH), (C) 5 mL trichloromethane (TCM). For each extraction, tubes were sonicated for 10 minutes at room temperature, then rotated end-over-end for 2 hours. Tubes were then centrifuged for 10 minutes, and the liquid phase was transferred to a clean glass centrifuge tube. 5 mL of TCM and 5 mL of LC-MS grade water were added, then the sample was centrifuged for 10 minutes to induce phase separation. The upper (aqueous) phase was removed and discarded via pasteur pipette. The bottom (organic) layer was re-extracted with 5 mL of TCM, and 5 mL of LC-MS grade water and the aqueous phase removed again. The remaining organic phase was evaporated at 30°C under a stream of N_2_ in a TurboVap LV until approximately 2 mL of solvent remained. To remove residual water and polar impurities, the remaining organic phase was run through a drying column consisting of 200 mg 8:2 w/w silica sand:NaSO_4_ (anhydrous). The drying column was rinsed with 2X volumes of TCM and MeOH before the sample was allowed to drain into a collection vial and dried to completeness under N_2_ at 30°C.

Intact phospholipids were separated from other lipid classes using a silica-gel column. Each column was conditioned with 5 mL of acetone and 10 mL of TCM. The sample was resuspended in 2 mL of TCM and applied to the column. Neutral lipids were separated by adding 5 mL of TCM to the column. Glycolipids were separated by adding 5 mL of acetone to the column. Phospholipids were separated with 5 mL of MeOH and collected, then dried to completion at 30°C. Intact phospholipids were derivatized to fatty acid methyl esters (FAMEs) using a KOH-MeOH derivatization (61). Samples were resuspended in 0.5 mL TCM and 0.5 mL MeOH, followed by 1 mL of methanolic KOH (KOH-MeOH; 0.45 g KOH dissolved in 40 mL anhydrous methanol). Samples were heated at 37°C for 30 minutes, then allowed to cool to room temperature. 2 mL of hexane and 0.2 mL of 1 M acetic acid were added to terminate the reaction. 2 mL of H_2_O was added and vortexed for 30 seconds to extract FAMEs from the organic layer; phase separation was induced by centrifuging for 1 minute. The top (organic) phase was transferred to a clean vial, and the sample was re-extracted again, before the FAME extract in hexane was dried under N_2_.

FAMEs were analyzed by GC-MS using a ZB-5ms (30 m, 0.25 mm ID, 1.00 µm film) column. Palmitic acid isobutyl ester (PAIBE) was applied as an external quantification standard. Samples were identified by comparing mass spectra against known standards and by comparison to BAME and 37-FAME (Supelco) chromatography standards.

FAME ^2^H/^1^H isotopic composition was determined using GC-P-IRMS: a Thermo Scientific Trace 1310 GC, pyrolysis reactor (1420°C) coupled to a Thermo Scientific 253 Plus IRMS via a Conflo IV interface. The GC was equipped with a ZB-5ms column (30 m, 0.25 mm ID, 1.00 µm film thickness). A C21 internal FAME standard was introduced into each sample run via co-injection.

Lipid-SIP soil water ^2^H/^1^H was determined by cavity ringdown spectroscopy (Picarro L2140i) against a standard curve of ^2^H-enriched standards. Each sample and standard was injected as 9 x 1 µL replicates to account for sample carryover; the first 3 replicates of each set were ignored. Highly ^2^H-enriched samples (from the vapor-SIP experiments) were gravimetrically diluted 1:100 with natural-abundance H_2_O before introduction to the instrument.

Lipid-SIP-derived growth rates (µ) were calculated using the relationship described by Caro et al. and Kopf et al. (23,62). In brief:

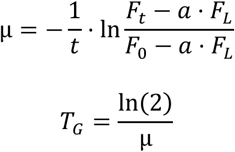

Where F_T_, F_L_, and F_0_ correspond to the ^2^H isotope fractional abundance of lipid biomass at time of sampling, isotope label, and initial lipid biomass, respectively.

### DNA Extraction and Metagenomic Sequencing Analysis

DNA was extracted in [biological] triplicate from soil using a Qiagen Powersoil Pro Kit following the manufacturer’s instructions. DNA was quantified on a QuBit fluorometer using 1X dsDNA HS mode. Libraries were prepared with a NEBNext Ultra II FS DNA Library Prep Kit. Libraries were sequenced on an Illumina NovaSeq X+.

Sequences were cleaned using the read_qc module of Metawrap (63). Contigs were assembled with MEGAHIT (64) using k-mer lengths 21, 33, 55, 77, 99, 127. MAGs were binned using semibin, metabat2, and metadecoder (65–67). Bins from these three tools were refined using the bin_refinement module of Metawrap. 73 individual MAGs in total were generated. MAG taxonomy was assigned with GTDB via gtdbtk (68).

Dereplicated MAG open reading frames (ORFs) were annotated with Bakta (v.1.11.4) (database v6.0, full) using default parameters (69). Minimum doubling times were calculated using codon usage bias statistics in R (v. 4.4.0) (70) via gRodon2 (40,41) in “partial” mode, which excludes pair-bias from the output. For nucleotide skew analyses, codon positions were classified by their degeneracy using the standard bacterial genetic code. Degeneracy is defined as the number of possible nucleotide substitutions that encode the same amino acid. Fourfold degenerate positions were treated as synonymous sites; onefold degenerate positions were treated as non-synonymous sites. The initiating Met and stop codon were excluded from all counts. GC and AT skew were calculated as:

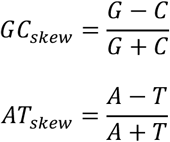

Where G, C, A, and T represent the counts of each nucleotide at relevant degenerate positions. Positions with ambiguous nucleotides are excluded. Functional categories were assigned to all annotated CDS using Clusters of Orthologous Genes (COG) categories parsed from the Bakta annotations. To characterize the distribution of synonymous skew across individual genes, GC/AT skew were computed separately for each annotated CDS. Genes with fewer than 10 fourfold-synonymous positions were excluded. Per-gene skew values were grouped by COG category and treatment condition. COG categories containing fewer than 15 qualifying genes in either condition were excluded from visualization.

### DNA Stable Isotope probing

DNA was extracted in triplicate from soil using a Qiagen Powersoil Pro Kit following the manufacturer’s instructions with one exception: during the final elution step, 100 µL of molecular grade water was substituted for Solution C6 (elution buffer), as Solution C6 contains compounds that interfere with ^18^O/^16^O analysis. DNA was quantified on a QuBit fluorometer in triplicate using 1X dsDNA HS mode. Sample DNA was mixed with sucrose (C_12_H_22_O_11_) to provide sufficient oxygen for IRMS analysis. Sucrose was prepared as a solution such that 194 µg of sucrose, or approximately 100 µg of sucrose O (sucrose is ∼51.4% O by weight), was added into a silver capsule as 30 µL of 6.485 µg/µL sucrose solution. 10 sucrose-only controls were added to establish a baseline isotopic value for mass balance calculations. Sucrose solutions were dried in silver capsules in a 96 well plate within a forced air incubator at 50°C for 24 hours. DNA extract was subsequently added to the capsules to achieve 0.75 µg of DNA oxygen per sample (DNA is ∼31.21% oxygen by weight). Samples were again allowed to dry for 24 hours, before silver capsules were sealed.

DNA oxygen isotopes were analyzed on an Elementar vario PYRO cube elemental analyzer interfaced to an Elementar precisION IRMS (Elementar Analysensysteme GmbH). Samples were thermally decomposed to CO in a glassy carbon reactor filled with glassy carbon, graphite felt, and lamp black at 1450°C. CO is isolated from N_2_ by an adsorption trap. Values for samples and standards are based on reference gas peaks introduced to the IRMS with each sample. Isotope variability is corrected by blank subtraction, regression of drift references, regression of peak area references, and normalization to bounding isotopic references. Lab references are calibrated against IAEA-V9, IAEA-600, USGS-42, USGS-43, USGS-35, and final delta values are expressed relative to VSMOW. Nylon, alanine, cellulose, chitin, caffeine (IAEA-600), keratin, sodium nitrate (USGS-35), and cellulose (IAEA-V9) were used as reference materials. Mean standard deviation for reference materials was ± 0.31 ‰. Total O introduced to the instrument was 114.8 ± 4.9 µg per sample.

For DNA-SIP calculations, DNA ^18^O values were converted from delta notation to fractional abundance (^18^O / (^18^O + ^17^O + ^16^O); the mean of triplicate samples calculated. ^18^O values were then adjusted by mass-balance to account for the dilution of DNA by sucrose O during preparation. Final values were adjusted to account for the presence of necromass-derived DNA, assuming that roughly 40% of DNA in soil is derived from extracellular (non-living) sources, as estimated by Carini et al. (71,72). DNA-SIP-derived growth rates were calculated using the same relationship as with lipid-SIP using ^18^O fractional abundance values, where F_L_ is the applied isotope label (corrected for the contribution of soil-derived water), F_T_ is the ^18^O isotopic enrichment at time of sampling, F_0_ is ^18^O isotopic enrichment at the start of the experiment, and a_w_ is 0.33, as reported by Hungate et al. (24).

### Targeted Metabolomics

Polar metabolites were extracted in triplicate following the method defined by Swenson & Northern (73). First, 8 mL of LC-MS grade water was added to 2 g of soil. Samples were sonicated at 50% amplitude for 2 x 30 seconds to release intracellular metabolites, then placed on a sample rotator at 30 rpm for 1 hour. Slurries were centrifuged at 3220G for 15 minutes at 4°C, then filtered using a 0.22 µm PES syringe filter. Samples were then frozen at-80°C then lyophilized for 24 hours. Dried extracts were held at-20°C prior to GC-TOF-MS and HILIC-LC-QTOF-MS analyses.

GC-TOF-MS analysis was conducted on a Gerstel ALEX-CIS GC-TOF-MS equipped with an RTX-5Sil MS (30 m x 0.25 mm x 0.25 µm film, 95% dimethyl, 5%diphenylpolysiloxane) column (Restek) using He as the carrier gas with a flow rate of 1 mL min^-1^. Sample was injected as 0.5 µL into a split-splitless inlet with a multi-baffled glass liner, with 25 splitless time. The injection temperature ramped from 50°C to 250°C by 12°C s^-1^. Oven temperature was held at 50°C for 1 min, then ramped at 20°C min^-1^ to 330°C, then held constant for 5 min. A Leco Pegasus IV MS was used with mass from 80 – 500 Da collected at 17 Hz and unit mass resolution; electron-impact ionization at-70 eV and 1800 V detector voltage. The transfer line and ion source were held at 230°C and 250°C, respectively. Inlet liners were exchanged after each set of 10 injections to reduce sample carryover.

Raw data files are preprocessed with ChromaTOF v2.32: no smoothing, 3 second peak width, baseline subtraction, and automatic MS deconvolution and peak detection using signal:noise of 5:1. Resulting text files are exported for additional filtering using the a BinBase algorithm as follows: validity of chromatogram (<10 peaks with intensity >10^7^ cps), unbiased retention index marker detection (MS similarity>800, validity of intensity range for high m/z marker ions), retention index calculation by 5^th^ order polynomial regression. Spectra are cut to 5% base peak abundance and matched to database entries from most to least abundant spectra using filters: retention index window +/-2 s retention time, validation of unique ions and apex masses (unique ions must be included in apexing masses and present at >3% of base peak abundance), MS similarity must fit criteria of peak purity and SNR and a final isomer filter. Quantification is reported as peak height using the unique ion as default.

HILIC-LC-QTOF-MS was conducted on a Waters Acquity Premier UPLC connected to a Sciex TripleTOF 6600 mass spectrometer. A BEH Amide column (1.7 µm, 2.1 mm x 50 mm) was used at 45°C with two mobile phases: (A) ultrapure water with 10 mM ammonium formate + 0.125% formic acid, pH 3 (B) 95:5 v/v acetonitrile:ultrapure water w/ 10 mM ammonium formate + 0.125% formic acid, pH 3. The following gradient was applied: 0 min, 100% B; 0.5 min, 100% B; 1.95 min, 70% B; 2.55 min, 30% B; 3.15 min, 100% B; 3.8 min, 100% B. The flow rate was 0.8 mL/min, and an injection volume of 1 µL from a 100 µL acetonitrile/water suspension (with internal standard mix) was used. Chromatograms were quality checked against internal standards for consistency of peak height and retention time. Then, raw data files were processed using MS-DIAL software to align and annotate peaks using an in-house mzRT library and MS/MS spectral matching with NIST/MoNA libraries. All MS/MS annotations were manually reviewed to ensure only high-quality compound identifications are included in the dataset. We note that, for dipeptides and tripeptides, orientation cannot be determined, and that Leu/Ile cannot be distinguished.

For both GC-TOF-MS and HILIC-LC-QTOF-MS analyses, raw peak heights were vector-normalized to reduce the impact of within-sequence drifts of instrument sensitivity. The sum of all peak heights of all identified metabolites for each sample (mTIC) is calculated. Data is normalized to the total average mTIC as follows for metabolite *i* of sample *j*:

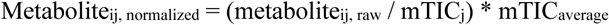

Where metabolite represents the peak height of a given metabolite, either ‘normalized*’* or ‘raw*’*, mTIC is the sum of peak heights for all annotated metabolites of sample *j* or ‘averaged’ across all datasets.

Shannon diversity (H) within a given sample is calculated as:

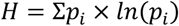

Where p_i_ is the molar proportion of the total metabolite pool comprised of metabolite *i*.

### Abiotic CO_2_ desorption experiment

Pure silica sand was combusted at 200 °C for 2 hours to remove microorganisms and organics. Sand was weighed into an acrylic cylinder. Deionized water, equilibrated with CO_2_ in laboratory air, was added to the experiment to simulate 20 mm of incident precipitation. Resulting CO_2_ concentrations at 10 cm depth in the sand were monitored by a Vaisala GMP252 CO2 probe for 24 hours. Concentrations measured with the CO_2_ probe were consistent with effluxes measured via LI-7810 CH_4_/CO_2_/H_2_O trace gas analyzer used in field experiments. Results from this experiment are displayed in **Fig. S4.**

## Data Availability

Analysis scripts, visualization scripts, and primary data generated in this study are available at https://github.com/tacaro/Drought-SIP. Raw metagenomic reads are available on the NCBI Sequence Read Archive (SRA) via BioProject accession PRJNA1444211.

## Acknowledgements

We thank Lucia Fuchslueger, Alberto Canarini, John Magyar, Woody Fischer, Grace Solini, Ashley Maloney, Ruby Fu, and Tanner Judd for valuable discussions during the development of this project. We thank field assistants Sarah Garzione and Cate Holmes. We thank John Magyar for assistance with site instrumentation and setup. We are grateful to Ann Close, Jonah Brees, Alana Tessman, and the staff of the White Buffalo Land Trust for providing access to Jalama Canyon Ranch and associated weather data. We thank Steve Alison and Moises Raymundo Perea-Vega for providing access to Loma Ridge. We thank Michael Mathuri for assistance with hydrogen isotope ratio mass spectrometry. We thank Nathan Dalleska for assistance with water isotope analyses. We thank Ruby Fu and Oliver Croft for assistance with abiotic CO_2_ release experiments. Metabolomics data were collected and processed at the UC Davis West Coast Metabolomics Center. DNA ^18^O analysis via TC/EA-IRMS was conducted at the UC Davis Stable Isotope Facility. We thank Igor Antoshechkin and the Millard and Muriel Jacobs Genetics and Genomics Laboratory at Caltech for assistance with DNA library preparation. This work was supported by The Linde Center for Global Environmental Science and the Linde Center for Science, Society, and Policy at Caltech. T. Caro was supported by the Foster & Coco Stanback postdoctoral fellowship at the California Institute of Technology.

J. Ibarra Arriaga was supported by the Center for Environmental Microbial Interactions at the California Institute of Technology. Figs. 1C and 1D were created in BioRender by T. Caro and are licensed under a CC-BY-4.0 (https://creativecommons.org/licenses/by/4.0/).

## Notes

### Competing Interest Statement

The authors have declared no competing interest.

