## Supplemental Information for "SIP-enabled multi-omics reveals soil microbiome responses to drought and rehydration"

1. Division of Geological and Planetary Sciences, California Institute of Technology
2. Division of Biology and Biological Engineering, California Institute of Technology
3. White Buffalo Land Trust
4. Division of Engineering and Applied Science, California Institute of Technology

***Corresponding authors:**

Tristan Caro
Alex Sessions
Smruthi Karthikeyan

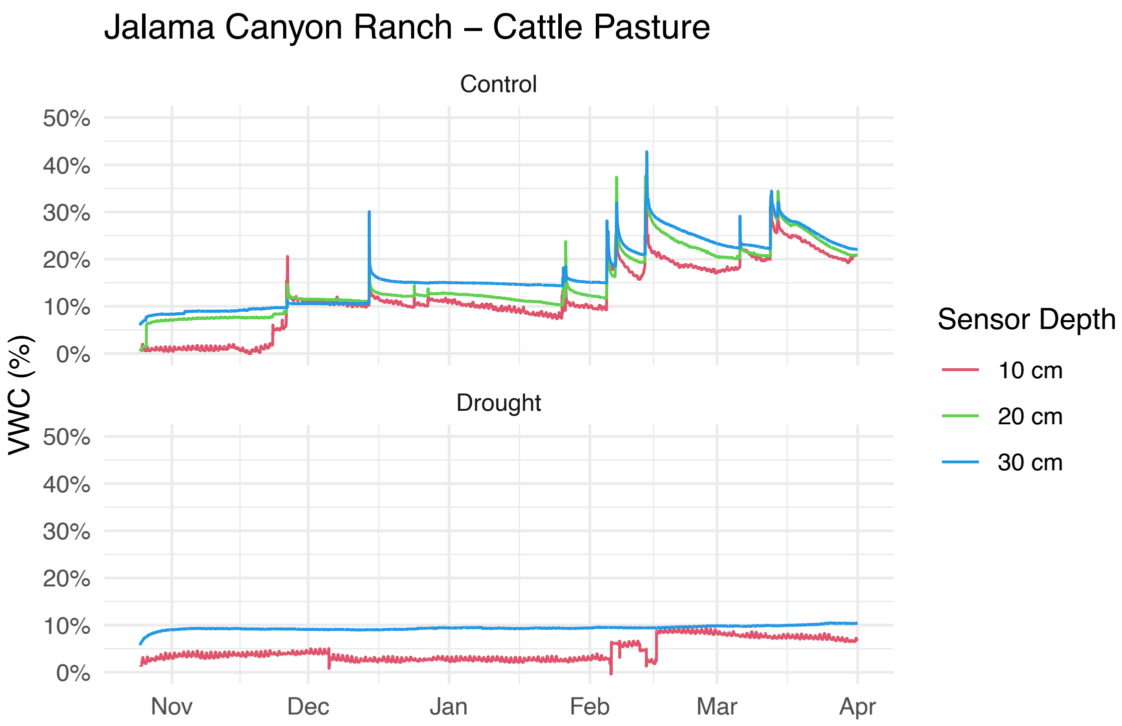


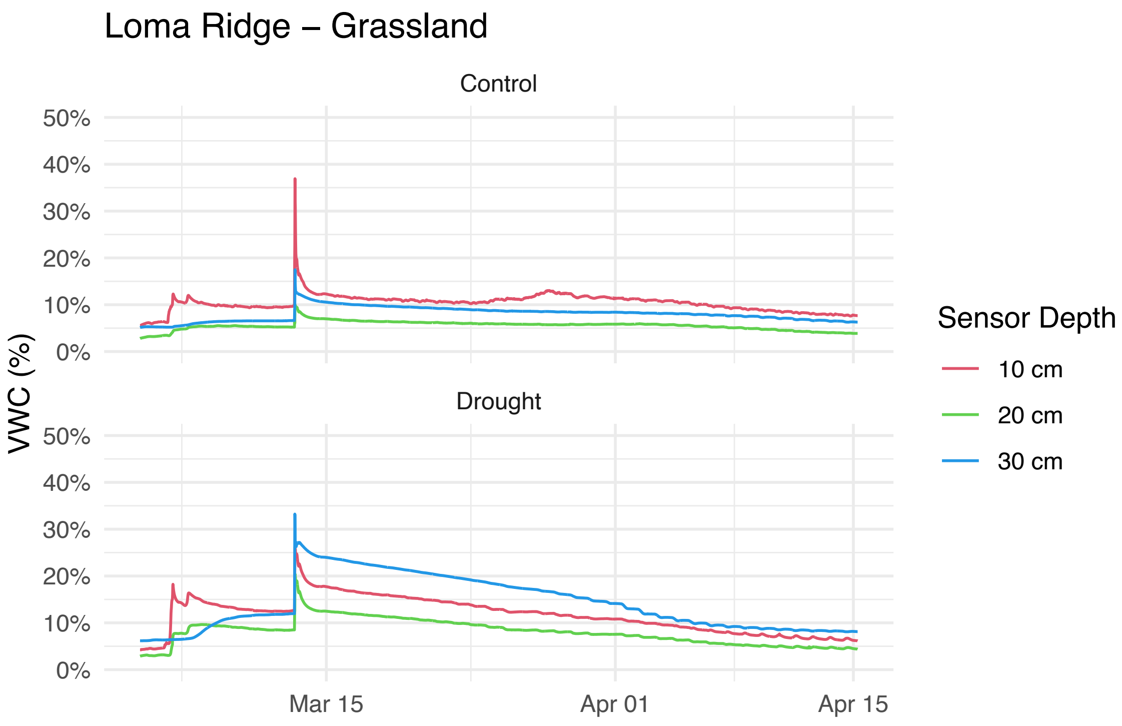


**Figure S1. Soil volumetric water content (VWC) across the experimental drought periods.**

Three sensors were deployed at each site. The 20 cm sensor at the Jalama Canyon Ranch drought plot failed to report accurate data.

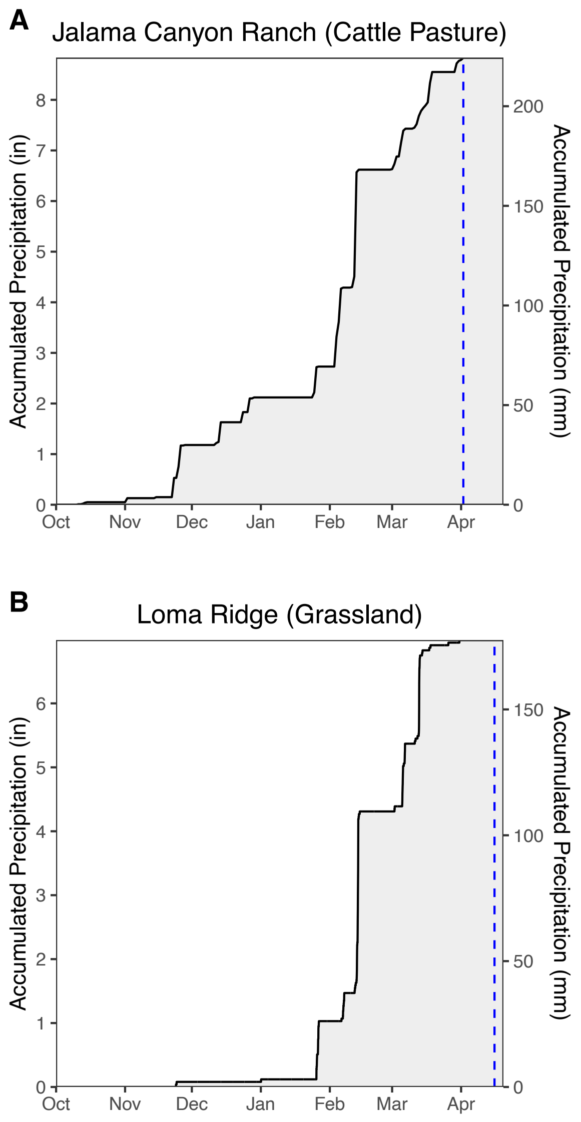


**Fig. S2.** Observed rainfall at Jalama Canyon Ranch (A) and Loma Ridge (B). In both plots, the dashed blue line indicates the time of soil sampling, gas flux monitoring, and SIP experiments. Data for Loma Ridge is accessed from the Hicks Canyon weather station dataset, maintained by Orange County Public Works http://hydstra.ocpublicworks.com/web.htm.

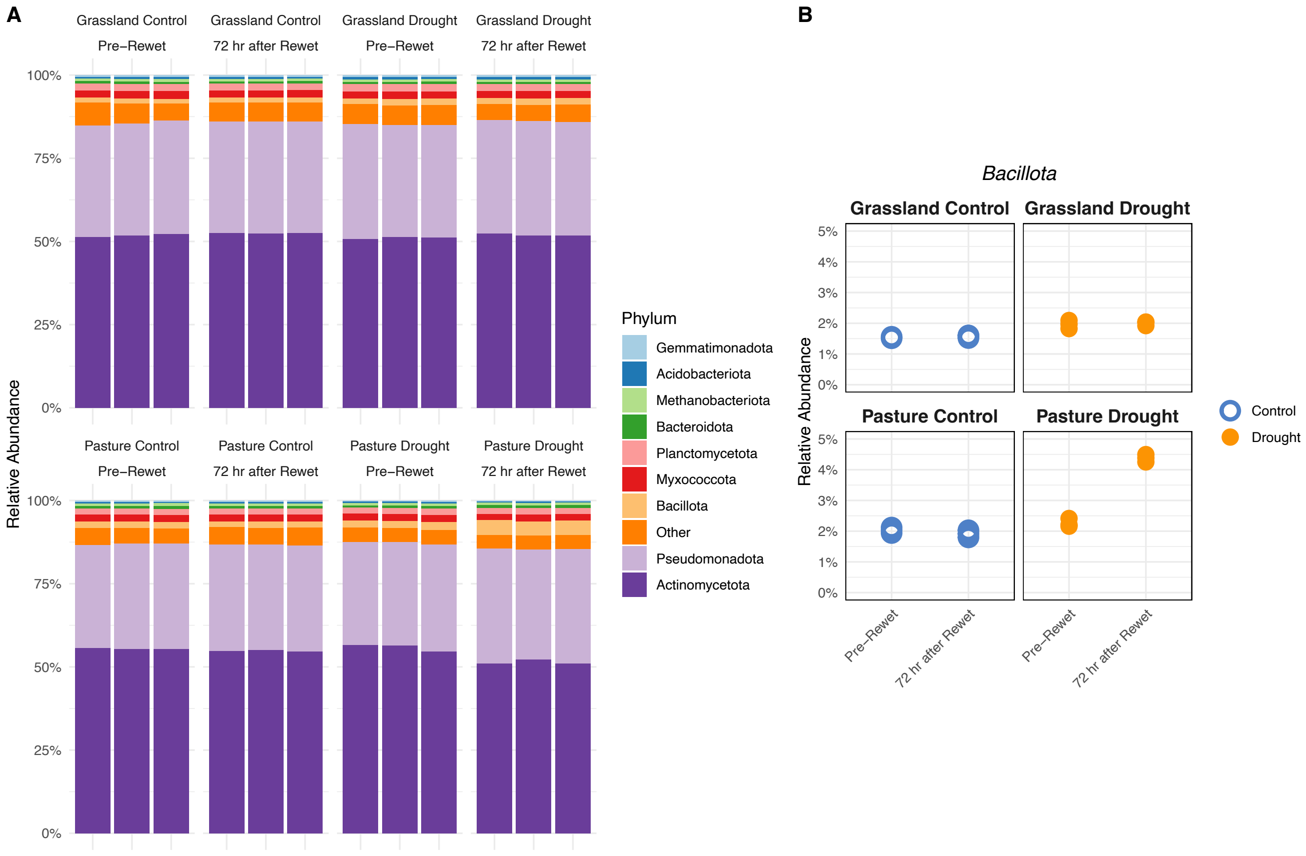


**Fig. S3. Read-based taxonomic abundance assessed via metagenomics.**
**(A)** Barplots indicating the relative abundance of phyla detected in our samples. **(B)** Relative abundance of *Bacillota* across treatments and time points.

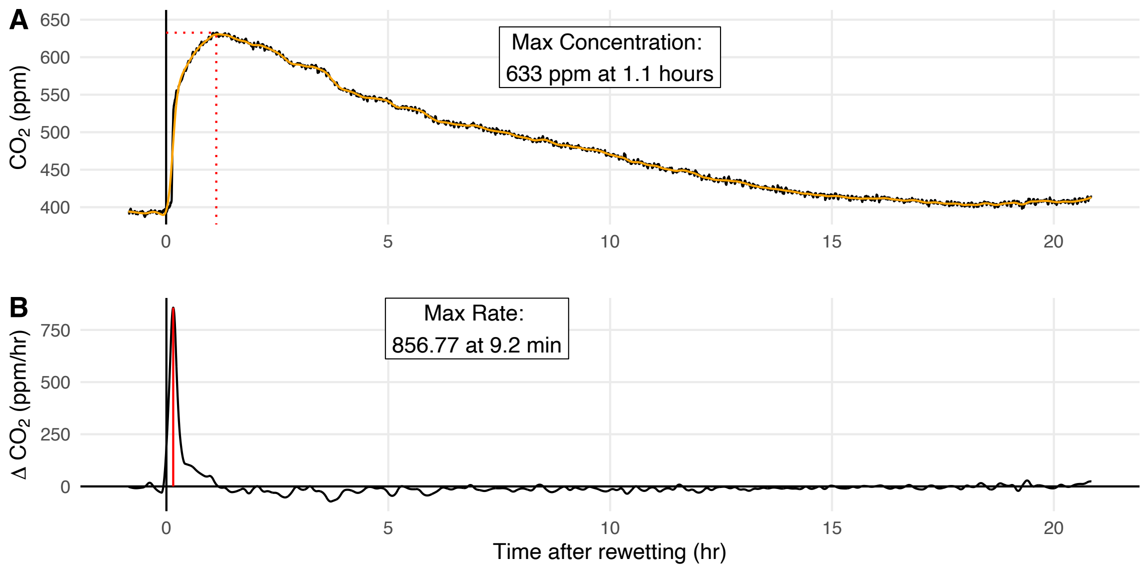


**Figure S4. Abiotic $CO_2$ release experiment.**

$CO_2$ concentration is displayed in **(A)**. The dotted red lines indicate the point at which maximum $CO_2$ concentration is observed. A smoothed spline (orange) is fit to the $CO_2$ data (black). **(B)** The first derivative of the smoothed spline is displayed, with the maximum noted with a vertical red line.

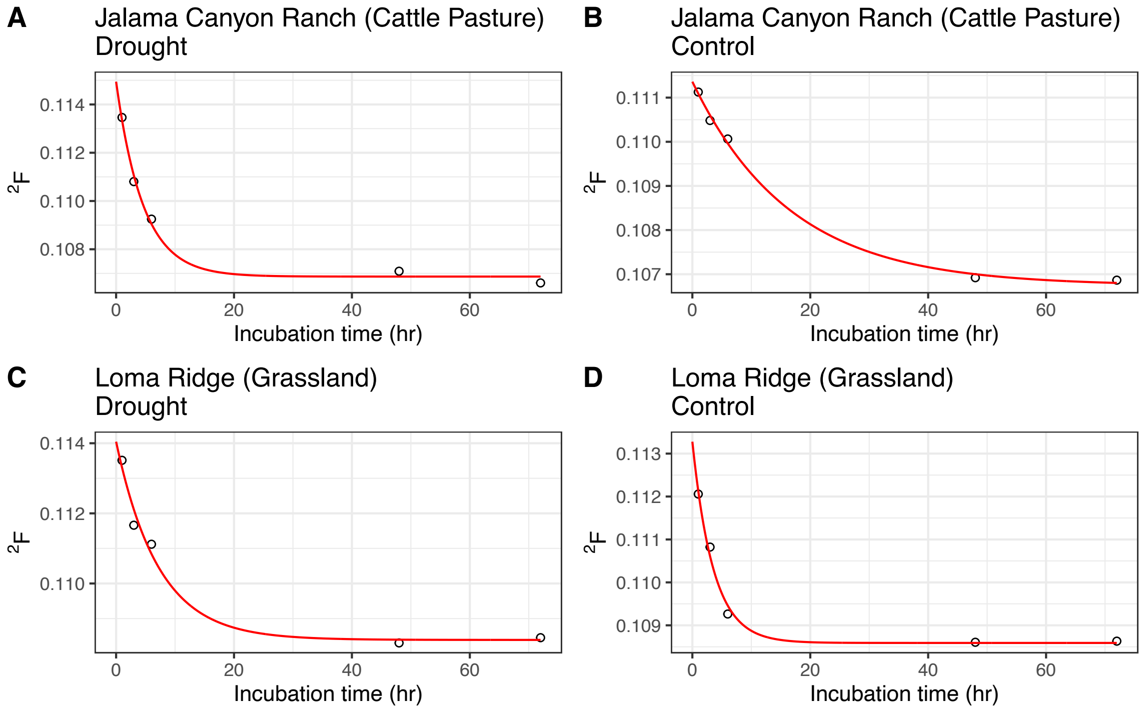


**Figure S5. Vapor-SIP equilibration curves.**

Hydrogen isotopic composition (as fractional abundance, 2F) as a function of incubation time. Red line indicates the best fit exponential decay function.

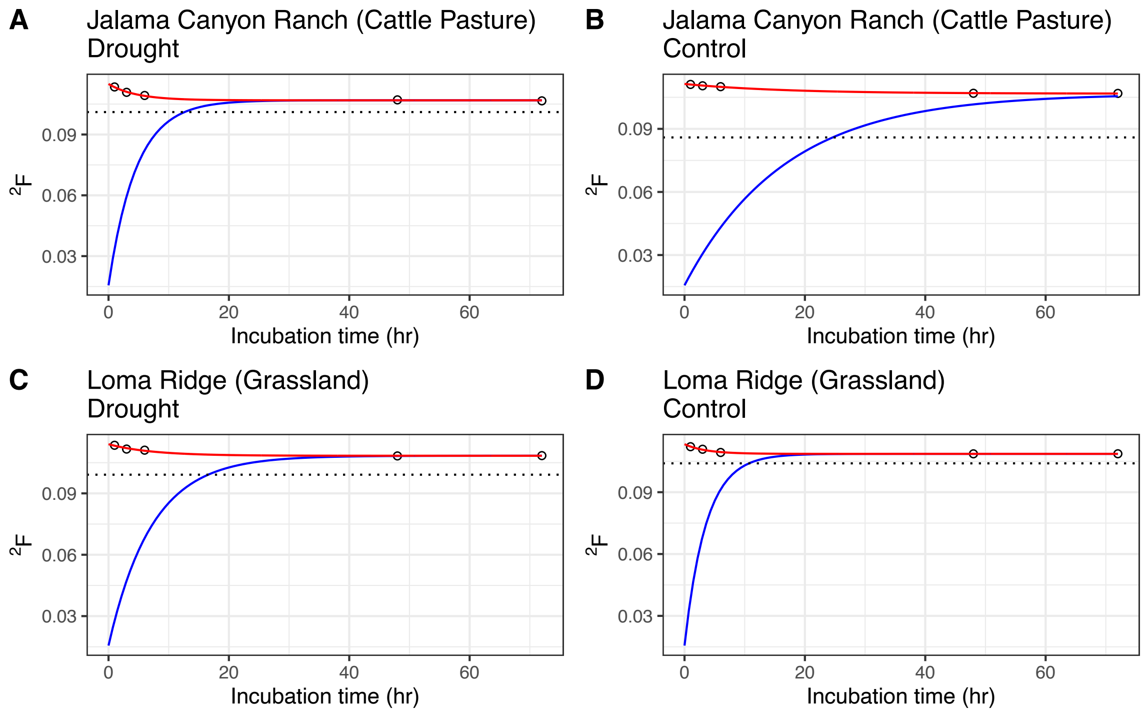


**Figure S6. Vapor-SIP equilibration curves.**

Blue line indicates the estimated soil porewater $^2$F as it equilibrates with the tracer solution.

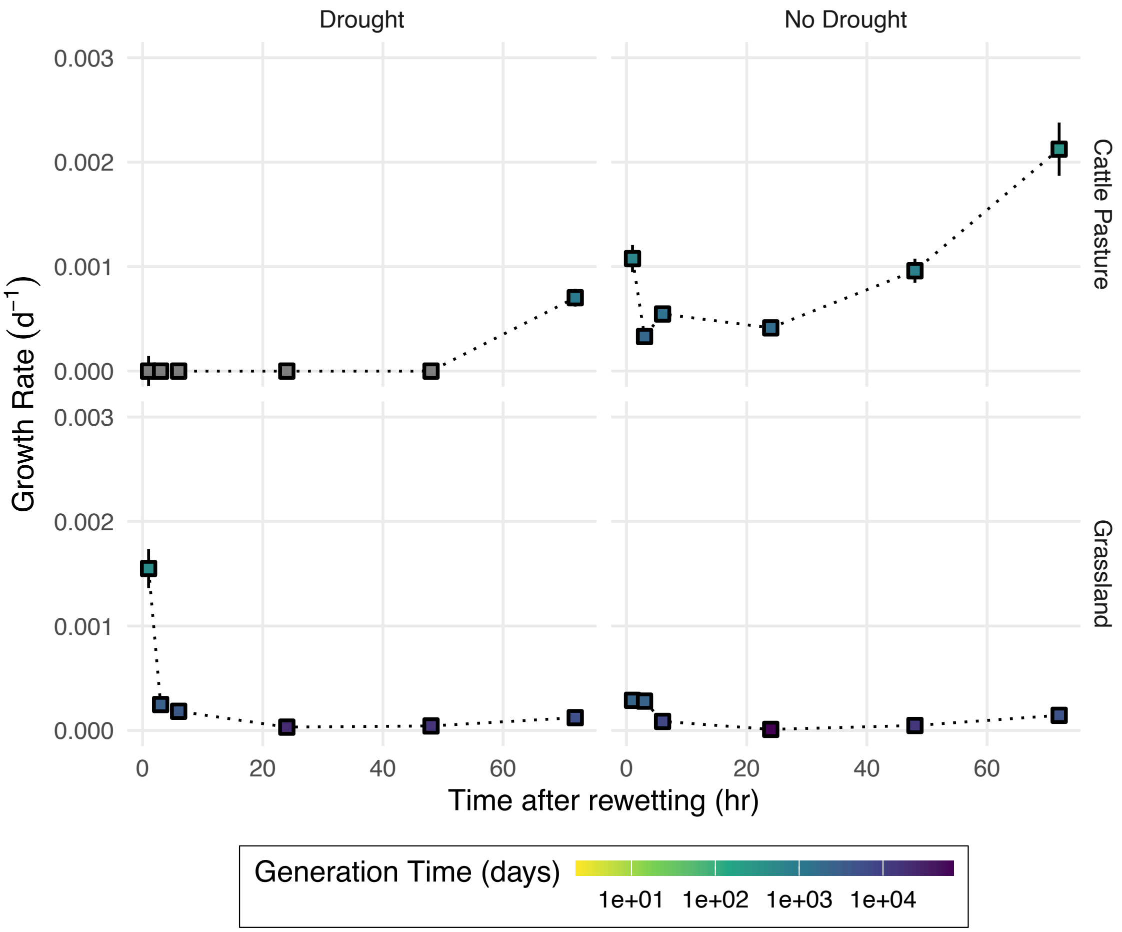


**Figure S7. Vapor-SIP growth rates.**

Growth rates from vapor-SIP experiments, identical to those in Fig. 2, but with the y-axis scaled to better illustrate differences along the time axis.

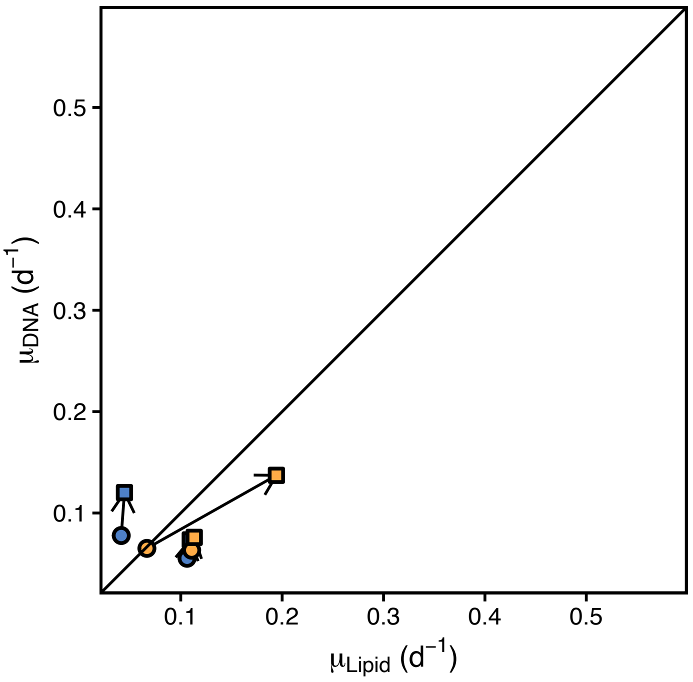


**Fig. S8.** Comparison of SIP techniques with DNA water assimilation efficiency and lipid water assimilation efficiency both held at 0.7. In other words, to achieve approximate parity, both lipid and DNA synthesis in soil acquires ~70% of H and O from water, respectively.

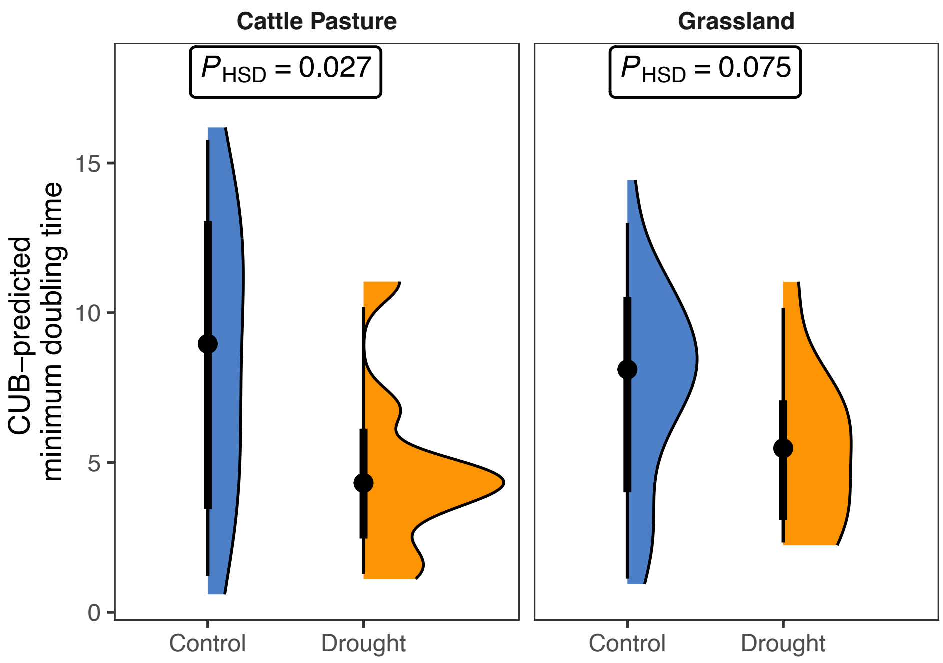


**Figure S9. CUB-predicted minimum doubling times, at each field site.**

Points, thick, and thin bars represent median, 66% CI, 95% CI, respectively. Significance indicated is Tukey's honestly significant difference test.

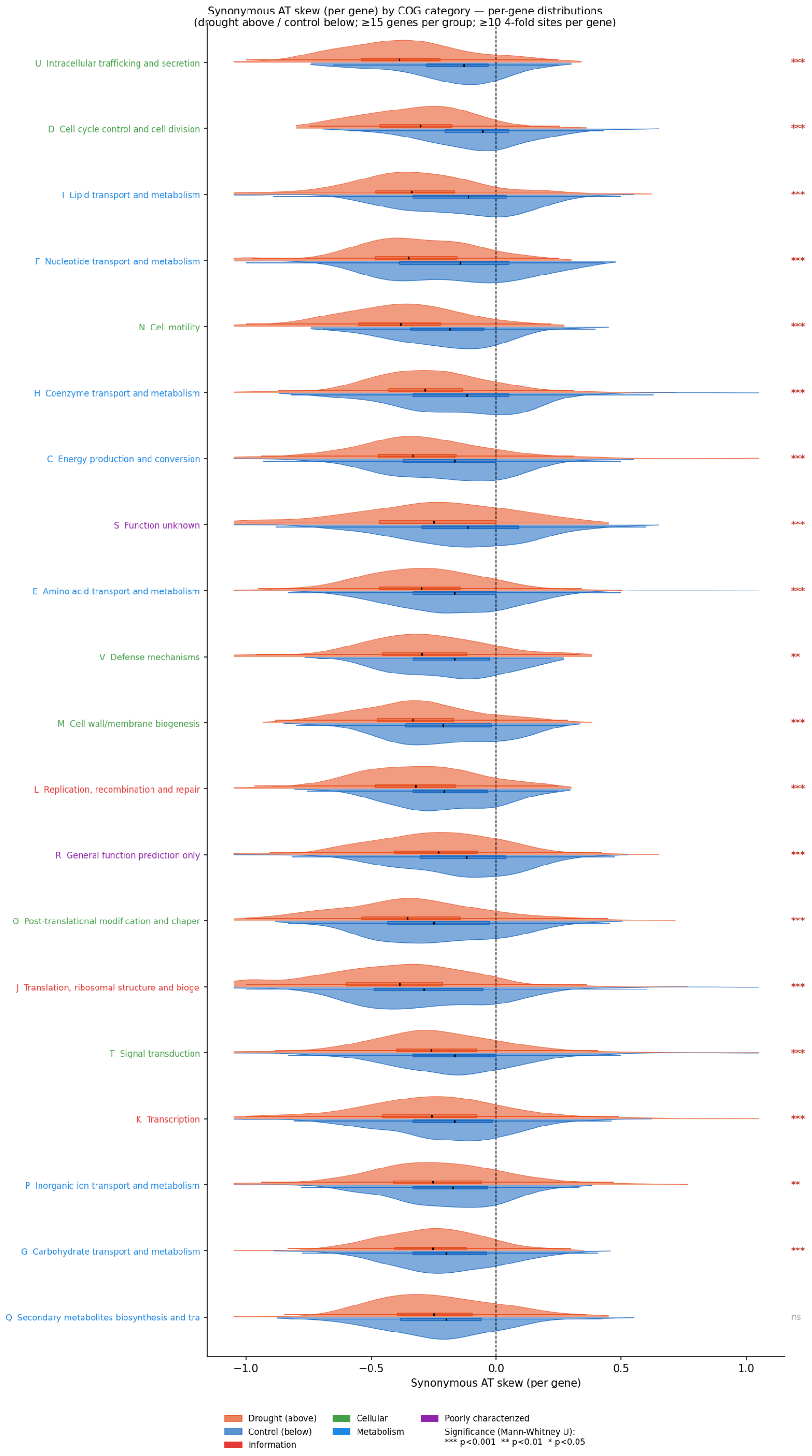


**Fig. S10.** Nucleotide skew (synonymous, AT) of genes annotated in MAGs from drought and control soils. COG categories are color-coded and indicated on the y axis. Results of Mann-Whitney U significance tests are displayed to the right of each comparison.

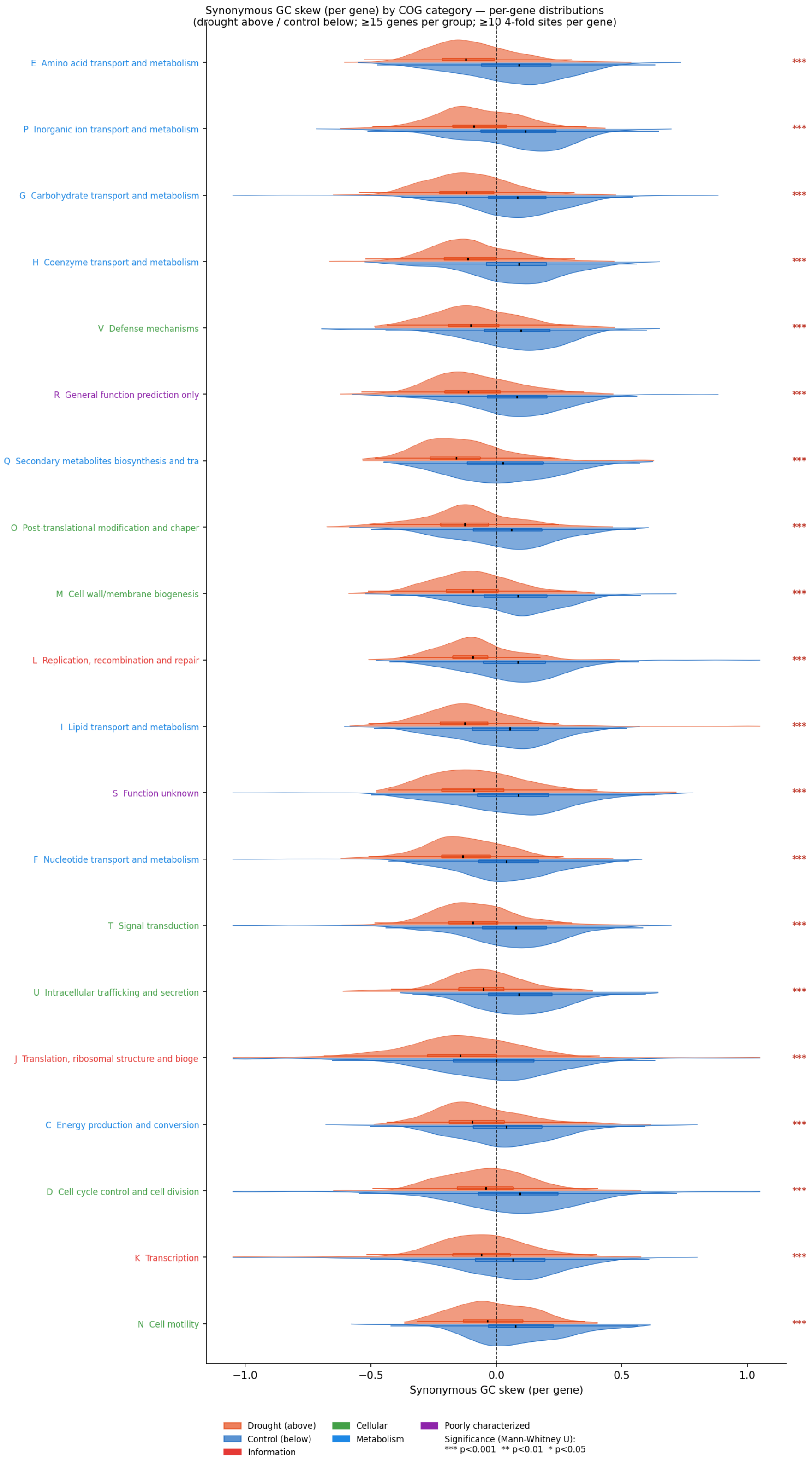


**Fig. S11.** Nucleotide skew (synonymous, AT) of genes annotated in MAGs from drought and control soils. COG categories are color-coded and indicated on the y axis. Results of Mann-Whitney U significance tests are displayed to the right of each comparison.

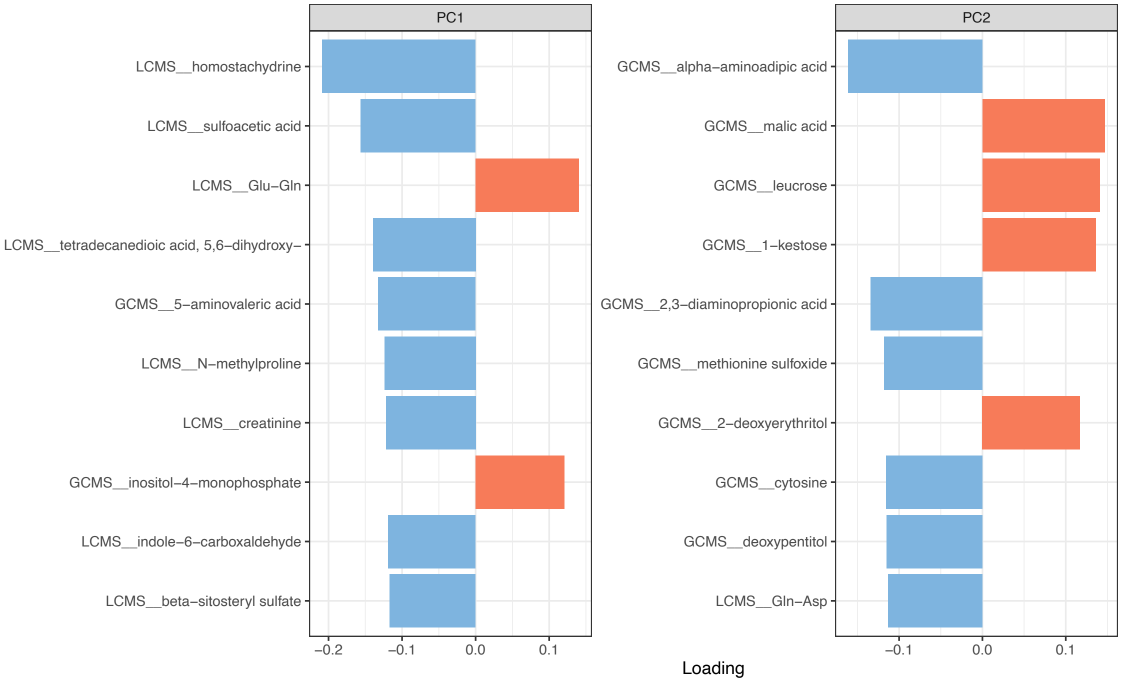


**Fig. S#.** Metabolomics dataset principal component (PC) loadings. Top 10 most significant loadings are displayed for each principal component. Principal components are plotted in **Fig. 5C**.
